# Rab29 GTPase is a specific cargo transport receptor to enhance COPII recruitment and anterograde trafficking of nascent GPCRs at ER

**DOI:** 10.64898/2026.11.23.768579

**Authors:** Xin Xu, Wei Huang, Mitsunori Fukuda, Nevin A. Lambert, Guangyu Wu

## Abstract

Rab GTPases are the master players in the cellular transport system, but their function and regulation in cargo recruitment to COPII vesicles at the endoplasmic reticulum (ER) are unknown. Here, we demonstrate that Rab29 inhibition via genetic deletion and inactivation markedly impedes the ER-to- Golgi transport, cell surface expression, and cellular functions of a subset of G protein-coupled receptors (GPCRs). More interestingly, Rab29 inhibition substantially attenuates GPCR concentration and recruitment kinetics at ER-derived COPII vesicles without affecting vesicle biogenesis. We also show that activated Rab29 localizes to the ER and COPII vesicles where it directly interacts with both its cargo GPCRs and all four Sec24 isoforms. We further identify a specific motif that mediates the interaction of angiotensin II type 2 receptor (AT2R) with Rab29; mutating this motif effectively blocks AT2R recruitment into COPII vesicles and compromises its delivery to the cell surface. Collectively, these data indicate that Rab29 is the first Rab functioning as a specific cargo transport receptor to enhance COPII capture and anterograde trafficking of GPCRs from the ER which is mediated through activation-dependent interactions with both Sec24 and cargo GPCRs and identify a novel function for the Rab family in membrane trafficking.

## Introduction

Vesicle-mediated molecular traffic between membrane-enclosed subcellular compartments is a fundamental process in eukaryotic cells. In the early secretary pathway, coat protein complex II (COPII) vesicles drive the selective delivery of newly synthesized secretory and membrane proteins, as well as lipids, from the endoplasmic reticulum (ER) to the Golgi complex. COPII vesicle formation initiates at ER exit sites (ERES) through Sar1 activation by the guanine nucleotide exchange factor (GEF) Sec12, triggering the subsequent recruitment of the heterodimeric Sec23-Sec24 and Sec13-Sec31 complexes (Downes and Zanetti, 2025; Farhan et al., 2025; Fath et al., 2007). For membrane cargo proteins, recognition, and concentration into COPII vesicles are primarily dictated by direct interactions with Sec24. Human cells express four Sec24 isoforms (A, B, C and D) that possess multiple distinct cargo- binding sites to accommodate a diverse array of client molecules (Miller et al., 2003). Meanwhile, soluble cargoes and certain membrane proteins lacking direct COPII-binding motifs rely on cargo- specific receptors, which interact simultaneously with COPII vesicles and cargo molecules, thereby bridging the cargo to the transport machinery. Several ER-localized transmembrane proteins have been identified in yeast and higher eukaryotes to function as cargo transport receptors (Barlowe and Helenius, 2016; Tang and Ginsburg, 2023).

Rab GTPases are a larger family of small G proteins that act as master regulators of vesicle- mediated membrane traffic. Mammals express more than 60 Rab proteins, each displaying distinct subcellular localizations and specialized functions in different cellular contexts and directing specific transport steps. As with all GTPases, the function of Rab GTPases are regulated by their cycle between an inactive GDP-bound state and an active GTP-bound state which are under control by guanine nucleotide exchange factors (GEFs) and GTPase-activating proteins (GAPs). In inactive state, Rab GTPases are extracted into the cytosol and bound by guanine nucleotide inhibitors (GDIs), whereas in active state, they are associated with specific target membrane and interact with effector proteins. At COPII-mediated ER export, Rab GTPases have been shown to orchestrate vesicle formation, motility, tethering, and membrane fusion with the Golgi apparatus (Carroll et al., 2001; Mizuno-Yamasaki et al., 2012; Stenmark, 2009). However, whether Rab GTPases can directly control cargo recruitment to COPII vesicles at the ER remains unknown.

Rab29 (also known as Rab7L1) is recognized as an atypical Rab because it binds nucleotides weakly and does not form complexes with GDIs in the cytosol (Gomez et al., 2019; Nagai-Ito et al., 2022), and its gene is encoded within the *PARK16* susceptibility locus and genetically linked to the pathogenesis of Parkinson’s disease (PD) (Beilina et al., 2014; Simón-Sánchez et al., 2009; Tucci et al., 2010). At the cellular level, Rab29 is predominantly localized to the Golgi and *trans*-Golgi network (TGN) (Wang et al., 2014) and is well known for recruiting and activating leucine-rich repeat kinase 2 (LRRK2) at the Golgi complex (Alessi and Pfeffer, 2024; Liu et al., 2018; Pfeffer, 2023; Purlyte et al., 2018; Steger et al., 2017; Vides et al., 2022; Zhu et al., 2023). It also regulates lysosomal homeostasis, lysosomal stress response (Eguchi et al., 2018; Komori et al., 2023; MacLeod et al., 2013), phagosome maturation (Pradhan et al., 2018), ciliogenesis (Onnis et al., 2015; Steger et al., 2017), retrograde protein transport from endosomes to the TGN (Beilina et al., 2020; MacLeod et al., 2013), and T-cell antigen receptor recycling (Onnis et al., 2015). However, its function and regulation at the ER are completely undefined.

As the largest family of membrane receptors, with over 800 members sharing a conserved structural topology, G protein-coupled receptors (GPCRs) regulate diverse cellular functions under both physiological and pathological conditions, making them the most intensively studied proteins for drug discovery (Kunselman et al., 2021; Liu et al., 2024; Lorente et al., 2025; Radoux-Mergault et al., 2023; von Zastrow and Sorkin, 2021). Like other plasma membrane (PM) proteins, GPCRs are synthesized, folded, and assembled in the ER, from which they are exported via COPII vesicles (Xu et al., 2023).

This ER export step critically dictates GPCR maturation kinetics, as well as receptor-elicited cellular response to hormones and drugs; its dysregulation causes various human diseases (Conn et al., 2007; Doly et al., 2016; Wei et al., 2019). While previous studies have identified numerous regulatory proteins and structural motifs that govern GPCR export from the ER (Assoumou et al., 2025; Sauvageau et al., 2014; Shiwarski et al., 2017; Xu et al., 2024a; Xu et al., 2024b), the exact mechanisms by which nascent GPCRs are recruited and packaged into COPII vesicles remain poorly understood.

Here, we demonstrate that Rab29 acts as a specific cargo transport receptor that promotes GPCR packaging into COPII vesicles, thereby facilitating their anterograde delivery from the ER. These findings reveal a novel function for the Rab family in membrane trafficking and identify an important mechanism governing COPII-directed GPCR forward trafficking.

## Results

### Rab29 is required for the ER-to-Golgi transport of specific nascent GPCRs

To systematically investigate the roles of the Rab family in the ER-to-Golgi transport of nascent GPCRs, we took advantage of a collection of MDCK knockout (KO) cells in which all mammalian *Rab* genes had been deleted individually or in combination (Homma et al., 2019), and measured the transport of nascent α_2A_- adrenergic receptor (α_2A_-AR), a prototypical GPCR, in the retention using selective hooks (RUSH) assays (Boncompain et al., 2012). In the RUSH system, cargo molecules were conjugated with enhanced green fluorescent protein (GFP) and a streptavidin-binding peptide (SBP), while an ER retention signal (KDEL) fused to streptavidin was used as a hook (Figure 1 – figure supplement 1). As expected, RUSH α_2A_-AR was colocalized with the ER marker Sec61β before incubation with biotin and was extensively colocalized with the ERGIC marker ERGIC53 and the Golgi markers giantin and β1,4- galactosyltransferase 1 (Gal T) after 30 min of biotin incubation in MDCK cells (Figure 1 – figure supplement 1). Therefore, for quantification, the concentration of α_2A_-AR (as well as other cargoes) in the juxtanuclear region after biotin induction was considered to transport to the Golgi complex, and ER- to-Golgi transport efficiencies were defined as the ratio of Golgi expression to total expression. After 10 min of biotin induction, α_2A_-AR was clearly concentrated at the Golgi, with the strongest Golgi expression observed at 30 min, after which the Golgi expression declined, indicating post-Golgi transport to the cell surface (Figure 1 – figure supplement 1).

Among the 58 Rabs studied, Rab26, Rab29 and Rab43 KO were found to markedly inhibit the transport of α_2A_-AR after 15 min of biotin incubation (Figure 1A, Figure 1 – figure supplement 2). This inhibition was observed at each time point of biotin induction in Rab29 KO cells (Figure 1B). In Rab1A, Rab1B and Rab2AB KO cells, α_2A_-AR was clearly exported from the ER but accumulated in punctate structures (Figure 1 – figure supplement 2), presumably due to the Golgi fragmentation as described previously (Homma et al., 2019). Therefore, we were unable to accurately quantify α_2A_-AR transport in these KO cells. In contrast, the ER-to-Golgi transport of α_2A_-AR was normal in other Rab KO MDCK cell lines as compared to control cells (Figure 1A, Figure 1 – figure supplement 2). Because the functions of Rab26 and Rab43 in the transport of GPCRs and other PM proteins along the secretory pathway have been reported previously (Haas et al., 2007; Li et al., 2012; Li et al., 2017), we focused on Rab29 whose functions at the ER are entirely unknown.

**Figure 1.**
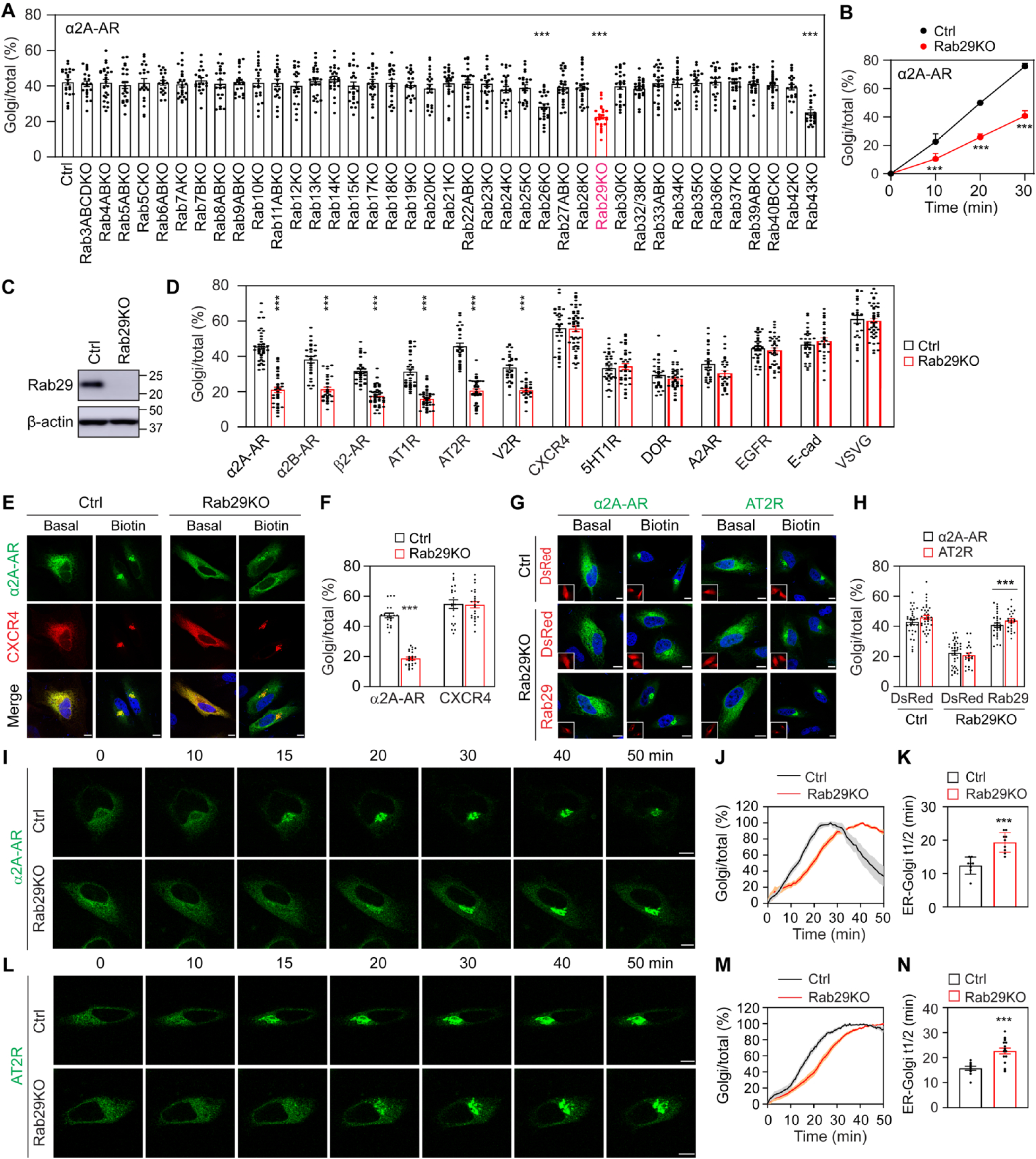
Screening for Rab29-regulated PM proteins using RUSH assays. **(A)** Screening for Rabs involved in the ER-to-Golgi transport of α_2A_-AR in MDCK cells. Parent and Rab KO cells were transfected with RUSH α_2A_-AR and then incubated with biotin for 15 min. ER-to-Golgi transport efficiencies were quantified as the ratio of Golgi expression to total expression. **(B)** The ER-Golgi transport of α_2A_-AR in parent and Rab29 KO MDCK cells after biotin induction for different time. **(C)** Rab29 expression in control and Rab29 KO HeLa cells. **(D)** Quantitative data showing the effects of Rab29 KO on the ER-to-Golgi transport of different PM proteins in HeLa cells. Cells were transfected with individual RUSH cargoes and then incubated with biotin for 15 min. **(E)** Effects of Rab29 KO on the ER-to-Golgi transport of GFP-α_2A_-AR and mCherry-CXCR4 co-expressed in the same HeLa cells. **(F)** Quantitative data shown in (E). **(G)** Rescue of the ER-to-Golgi transport of α_2A_-AR and AT2R by CRISPR-Cas9-resistant DsRed-Rab29 in Rab29 KO HeLa cells. Insets show the expression of DsRed or DsRed-Rab29. **H** Quantitative data shown in (G). **(I)** and **(L)** Live-cell imaging showing the effects of Rab29 KO on the export of α_2A_-AR (I) and AT2R (L). **(J)** and **(M)** Transport curves of α_2A_-AR (J) and AT2R (M) in control and Rab29 KO HeLa cells. Images were captured at 30-sec intervals. **(K)** and **(N)** t_1/2_ values of ER-to-Golgi transport of α_2A_-AR (K) and AT2R (N) in control and Rab29 KO cells. All quantitative data are presented as means ± SE; n = 20-26 cells in (A), 38-56 cells in (B), 26-52 cells in (D), 20-23 cells in (F), 20-35 cells in (H), 8-9 cells in (J) and (K), and 10-20 cells in (M) and (N), each from 3-5 different experiments. ***p < 0.001 versus respective control in (A), (B), (D), (F), (K) and (N), and versus Rab29 KO cells transfected with DsRed in (H). Scale bars,10 µm.

To further define the trafficking function of Rab29, we determined its role in the ER-to-Golgi transport of several different PM proteins in additional cell types. Accordingly, we generated Rab29 KO HeLa and HEK293 cells and also used the transient expression of the inactive GDP-bound mutant T21N (TN) and cytosolic mutant D63A (DA) (Gomez et al., 2019) to inhibit Rab29 function. The PM proteins studied included ten GPCRs [α_2A_-AR, α_2B_-AR, β_2_-AR, angiotensin II (Ang II) type 1 and 2 receptors (AT1R and AT2R), vasopressin V2 receptor (V2R), CXCR4, 5-hydroxytryptamine receptor 1B (5HT1R), δ opioid receptor (DOR), and adenosine A2A receptor (A2AR)] and three non-GPCR proteins [epidermal growth factor receptor (EGFR), E-cadherin and vesicular stomatitis virus glycoprotein (VSVG)]. In HeLa cells, the ER-to-Golgi transport of six GPCRs (α_2A_-AR, α_2B_-AR, β_2_-AR, AT1R, AT2R, and V2R) was markedly attenuated by 45-57% in Rab29 KO cells (Figure 1C and 1D, Figure 1 – figure supplement 2) and in Rab29 mutant-expressing cells (Figure 1 – figure supplement 3) as compared with control cells in RUSH assays after 15 min of biotin induction. By contrast, Rab29 KO and its inactive mutants did not affect the ER-to-Golgi transport of CXCR4, 5HT1R, DOR, A2AR, EGFR, E- cadherin, and VSVG in HeLa cells (Figure 1D; Figure 1 – figure supplement 3).

To confirm the differential effects of Rab29 KO on distinct GPCRs, RUSH α_2A_-AR and CXCR4 were co-expressed, and their transport was compared in cells expressing both receptors. α_2A_-AR and CXCR4 were similarly transported and extensively colocalized in control cells, whereas the ER-to-Golgi export of α_2A_-AR, but not CXCR4, was attenuated in Rab29 KO cells (Figure 1E and 1F). In rescue experiments, the expression of CRISPR-Cas9-resistant Rab29 almost completely restored the defective transport of α_2A_-AR and AT2R in Rab29-depleted cells (Figure 1G and 1H).

We then measured the effects of Rab29 inhibition on export kinetics using α_2A_-AR and AT2R as representative cargoes in live-cell RUSH assays. In control cells, both receptors were concentrated clearly at the Golgi approximately 10 min after the addition of biotin. The strongest Golgi expression of α_2A_-AR was observed at 20 to 30 min, and its Golgi expression began to decline after 30 min (Figure 1I and 1J). The strongest Golgi expression of AT2R was observed after 30 min of biotin incubation and remained elevated even after 60 min of induction (Figure 1L and 1M). These data suggest that α_2A_-AR and AT2R share similar ER-to-Golgi transport kinetics, but the post-Golgi transport of α_2A_-AR is much faster than that of AT2R. Compared with control cells, the time-course curves of α_2A_-AR and AT2R were clearly shifted to the right, with the half time (t_1/2_) values of ER-to-Golgi transport being significantly higher in Rab29 KO cells (Figure 1I-1N) and in Rab29TN-expressing cells (Figure 1 – figure supplement 4).

In HEK293 cells, Rab29 KO inhibited the ER-to-Golgi transport of α_2A_-AR and AT2R, but not VSVG (Figure 1 – figure supplement 5). The expression of CRISPR-Cas9-resistant Rab29 almost completely restored the defective transport of both α_2A_-AR and AT2R in Rab29 KO cells, whereas inactive Rab29 mutants neither rescued nor further exacerbated the transport defects (Figure 1 – figure supplement 5). Taken together, these data demonstrate that the normal function of Rab29 is required for the ER-to-Golgi transport of a specific subset of GPCRs after their synthesis in the ER.

### Rab29 regulates the surface expression and functions of GPCRs

To determine the effects of Rab29 on GPCR expression at the cell surface, the functional destination for most GPCRs, HiBiT (an 11 amino acid peptide) was inserted into the RUSH GPCR constructs between GFP and SBP. Surface receptor levels were then quantified by measuring the luminescence signal generated after addition of LgBiT (Figure 2A). After biotin induction for 60 min, the surface expression of α_2A_-AR, β_2_-AR, and AT2R was significantly decreased in Rab29 KO and inactive Rab29 mutant-expressing cells compared with their respective controls (Figure 2B). To study endogenous GPCRs, we knocked out Rab29 in human colon cancer HT29 cells (Figure 2C), which natively express α_2A_-AR and AT2R. As measured by radioligand binding of membrane preparations, surface α_2A_-AR levels were significantly reduced in Rab29 KO cells relative to control cells (Figure 2D).

**Figure 2.**
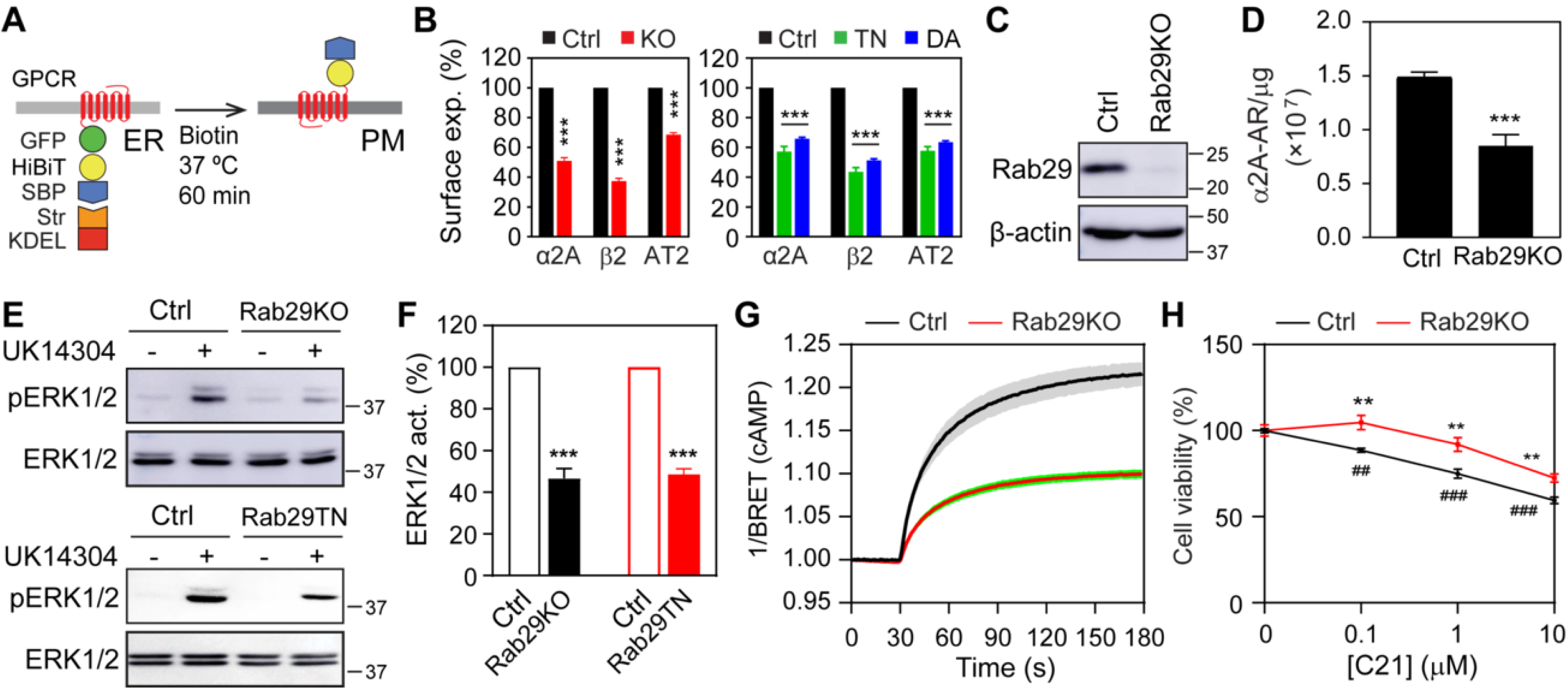
Rab29 KO and inactive mutants impede the cell surface expression and signaling of GPCRs. **(A)** Schematic diagram of the modified RUSH system used to quantify the cell surface expression of GPCRs in Nano-Glo HiBiT assays. **(B)** Rab29 KO (left panel) and its inactive mutants (right panel) inhibit the cell surface expression of GPCRs. To measure the effect of Rab29 mutants, HEK293 cells were co-transfected with GPCRs and Rab29 mutants. Surface receptors were quantified by measuring the luminescence signal following the addition of LgBiT and furimazine substrate. **(C)** Rab29 expression in control and Rab29 KO HT29 cells. **(D)** Effect of Rab29 KO on the cell surface expression of endogenous α_2A_-AR in HT29 cells, as measured by radioligand binding. Data are expressed as the number of α_2A_-AR per microgram of membrane protein. **(E)** Effect of Rab29 KO and expression of the Rab29TN mutant on α_2A_-AR-mediatd ERK1/2 activation. **(F)** Quantitative data shown in (E). **(G)** Effect of Rab29 KO on β_2_-AR-stimulated cAMP production. **(H)** Effect of Rab29 KO on HT29 cell proliferation in response to C21 stimulation. All quantitative data are presented as means ± SE; n = 3-4 independent experiments. **p < 0.01 and ***p < 0.001 versus the respective control; ^##^p < 0.01 and ^###^p < 0.001 versus control cells without C21. Str, streptavidin; α2A, α_2A_-AR; β2, β_2_-AR; AT2, AT2R.

To address whether Rab29 affects the ultimate function of GPCRs, we used ERK1/2 activation by α_2A_-AR, cAMP production by β_2_-AR, and the inhibition of cell proliferation by AT2R as readouts. ERK1/2 activation after stimulation with the specific α_2_-AR agonist UK14304 (Figure 2E and 2F) and cAMP production induced by the β_2_-AR agonist isoproterenol (ISO) (Figure 2G) were much lower in Rab29 KO HEK293 cells than in control cells. In addition, the expression of the Rab29TN mutant also inhibited ERK1/2 activation upon UK14304 stimulation in α_2A_-AR-transfected HEK293 cells (Figure 2E and 2F). Finally, in both control and Rab29 KO HT29 cells, treatment with the specific non-peptide AT2R agonist C21 dose-dependently inhibited cell proliferation (Figure 2H). However, this anti- proliferative effect was significantly attenuated in Rab29 KO cells across all tested C21 concentrations (Figure 2H).

### Rab29 controls the recruitment of selective GPCRs to COPII vesicles

To define how Rab29 regulates ER-to-Golgi transport, we examined the effects of Rab29 inhibition on the Golgi structural integrity, COPII vesicle formation, and cargo recruitment. Immunofluorescence after staining with antibodies against GM130 and p230 revealed that Rab29 KO did not overtly disrupt overall Golgi morphology (Figure 3 – figure supplement 1). Furthermore, the average numbers of COPII vesicles using GFP-Sec24B as a marker, as well as the signal intensity of GFP-Sec24B per vesicle, were similar among control cells, Rab29 KO cells and cells expressing Rab29 mutants (Figure 3 – figure supplement 1). In contrast, vesicle formation was almost completely abolished by the expression of the Sar1H79G mutant (Figure 3 – figure supplement 1).

To study cargo recruitment to COPII vesicles, we focused on AT2R, CXCR4, and VSVG, which are known to be highly concentrated in COPII vesicles (Xu et al., 2023). In the absence of biotin, RUSH AT2R, CXCR4, and VSVG accumulated in punctate structures, and nearly 100% of these cargo- carrying puncta contained the COPII marker Sec24B (Figure 3 – figure supplement 2). These cargo- containing puncta were markedly depleted by Sar1H79G (Figure 3 – figure supplement 2), confirming that they represent *bona fide* COPII vesicles. AT2R concentration within COPII vesicles significantly decreased in Rab29 KO cells (Figure 3A and 3B) and in cells expressing Rab29 mutants (Figure 3C and 3D) compared with control cells. In contrast, the recruitment of CXCR4 and VSVG to COPII vesicles remained unaffected by Rab29 KO or mutants (Figure 3A-3D).

**Figure 3.**
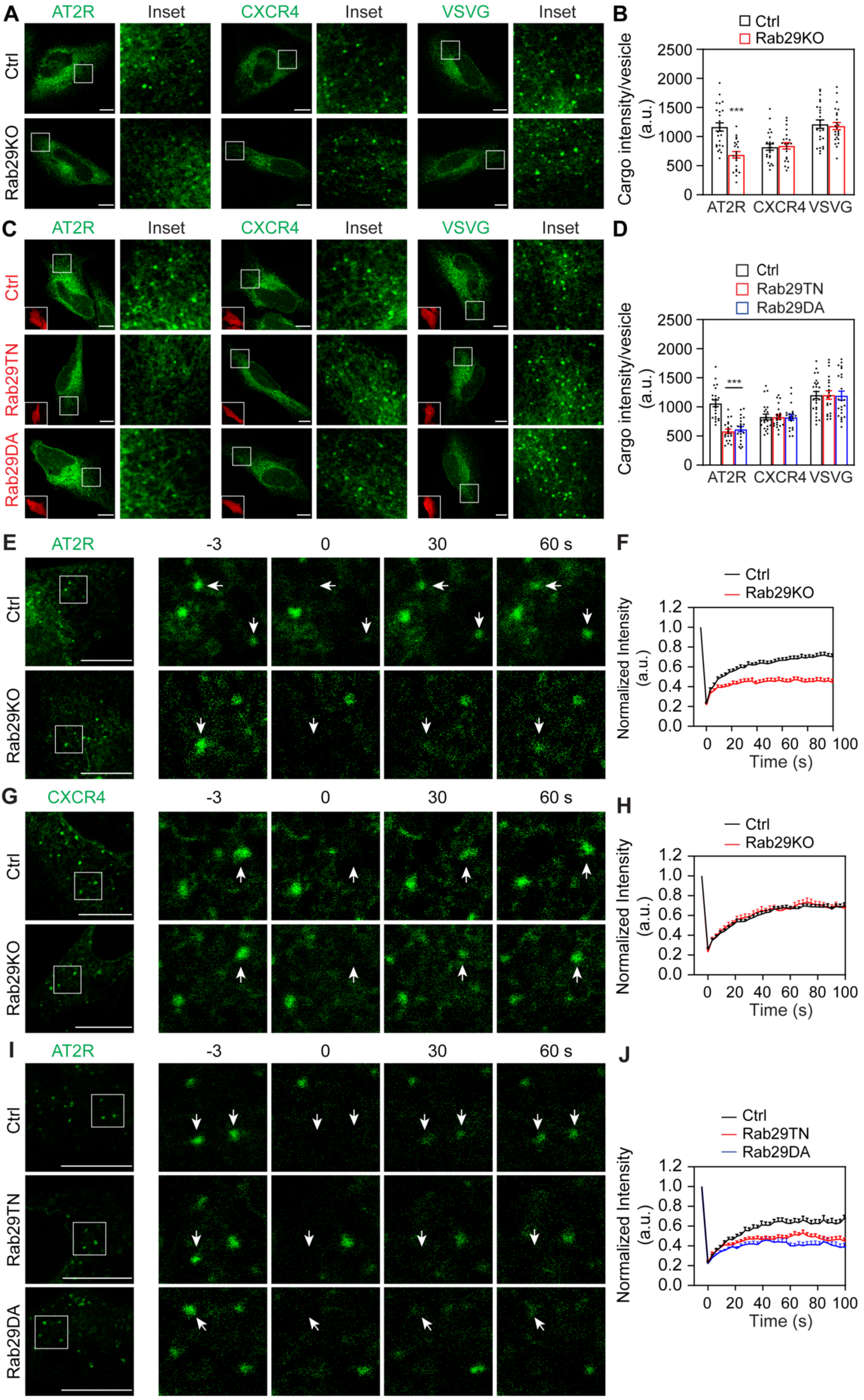
Rab29 KO and inactive mutants attenuate COPII recruitment of selective GPCRs. **(A)** and **(C)** Rab29 KO (A) and Rab29 mutants (C) inhibit the recruitment of AT2R, but not CXCR4 and VSVG, to COPII vesicles. **(B)** and **(D)** Quantitative data shown in (A) and (C), respectively. **(E)** and **(G)** Representative images showing the effects of Rab29 KO on the fluorescence recovery after photobleaching of AT2R (E) and CXCR4 (G) at COPII vesicles in HeLa cells. **(F)** and **(H)** Fluorescence recovery curves of AT2R (F) and CXCR4 (H). Images were captured at 3-sec intervals following photobleaching. **(I)** Recruitment kinetics of AT2R to COPII vesicles in cells expressing inactive Rab29 mutants. **(J)** Quantitative data shown in (I). Insets on the bottom left in (C) show the expression of DsRed or DsRed-Rab29 mutants. Arrows indicate the photobleached vesicles. All quantitative data are presented as means ± SE; n = 20-25 cells in (B) and (D), and 15-40 vesicles from 12-25 cells in (F), (H), and (J) from at least 4 separate experiments. ***p < 0.001 versus control. Scale bars,10 µm.

Fluorescence recovery after photobleaching (FRAP) assays were then employed to compare the COPII recruitment kinetics of AT2R and CXCR4 in live cells. Following photobleaching, the fluorescence intensities of both AT2R and CXCR4 at COPII vesicles gradually recovered but failed to return to prebleach levels in control cells (Figure 3E-3J). This incomplete recovery indicates a substantial fraction of immobilized receptors at the vesicles. AT2R recovery was markedly slower in Rab29 KO cells than in control cells (Figure 3E and 3F). In contrast, the recovery kinetics of CXCR4 remained virtually identical between Rab29 KO and control cells (Figure 3G and 3H). Consistent with the data obtained from Rab29 KO cells, expression of inactive Rab29 mutants also significantly delayed AT2R recovery (Figure 3I and 3J). Together, these data demonstrate that Rab29 selectively accelerates the COPII recruitment kinetics of AT2R, without influencing the biogenesis of the COPII vesicles themselves.

We next sought to determine the role of Rab29 in the COPII recruitment of α_2A_-AR. Although our previous studies suggest that the ER export of α_2A_-AR is mediated by COPII vesicles (Xu et al., 2023), we could not reliably detect the receptor within COPII vesicles using standard imaging or *in vitro* vesicle budding assays (data not shown), likely due to its low steady-state expression levels. To overcome this limitation, we biochemically evaluated its proximity to the COPII machinery using proximity-dependent biotinylation assays, in which either α_2A_-AR or Sar1B was conjugated to the promiscuous biotin ligase BirA R118G (BirA*) that catalyzes the biotinylation of neighboring proteins. This proximity assay is a well-established approach for identifying cargoes, cargo receptors, and transport machinery components within COPII-mediated transport pathways (Emmer et al., 2018; Nie et al., 2018; Qiu et al., 2023; Xu et al., 2024a). We found that the COPII components Sec24B and Sec24C were strongly biotinylated in cells expressing α_2A_-AR-BirA* (Figure 4A). Similarly, Sec24B and Sec24C were biotinylated in cells expressing AT2R-BirA* (Figure 4B). Under the same experimental conditions, both transiently expressed Rab29 (Figure 4C) and endogenous Rab29 (Figure 4D) were robustly biotinylated by α_2A_-AR-BirA*. Collectively, these data suggest that α_2A_-AR is in close spatial proximity to both the COPII component Sec24 and its regulator, Rab29.

**Figure 4.**
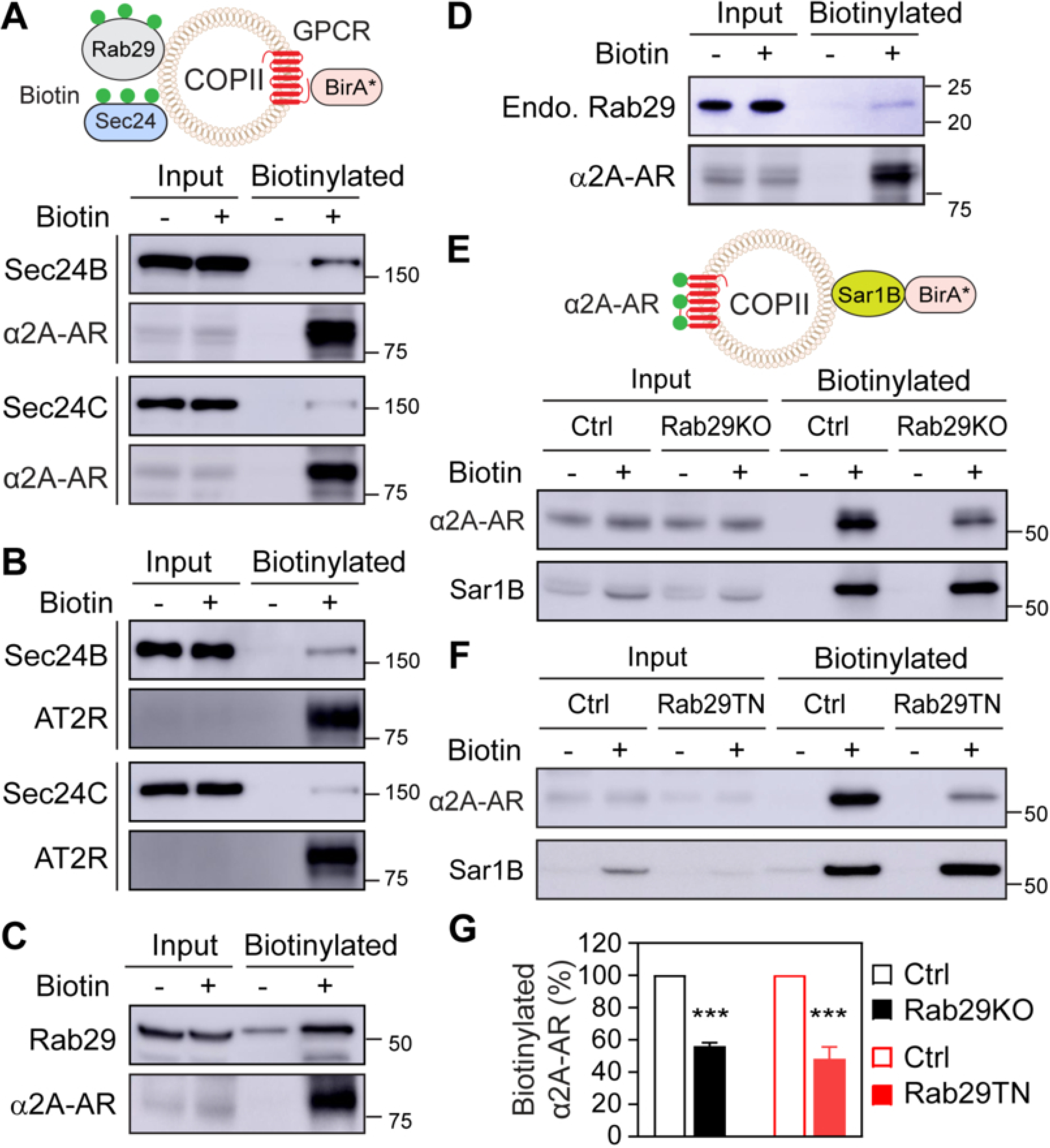
Rab29 inhibition attenuates α_2A_-AR biotinylation by Sar1B-BirA*. **(A)** and **(B)** Biotinylation of Sec24B and Sec24C by α_2A_-AR-BirA* (A) and AT2R-BirA* (B). HEK293 cells were transfected with α_2A_-AR-BirA*-HA (A) or AT2R-BirA*-HA (B), along with GFP-Sec24, and then treated with (+) or without (-) biotin for 24 h. Following pulldown with streptavidin-conjugated beads, Sec24 biotinylation was analyzed by immunoblotting using anti-GFP antibodies, whereas α_2A_-AR-BirA*-HA was detected by anti-HA antibodies. **(C)** and **(D)** Biotinylation of GFP-Rab29 (C) and endogenous Rab29 (D) by α_2A_-AR-BirA*. **(E)** Rab29 KO inhibits α_2A_-AR biotinylation by Sar1B-BirA*. Control and Rab29KO HEK293 cells were transfected with HA-α_2A_-AR and Myc-Sar1B-BirA*, and then treated with biotin for 24 h. Biotinylated α_2A_-AR and total Myc-Sar1B-BirA* levels were determined using anti-HA and anti-Myc antibodies, respectively. **(F)** Inhibition of Sar1B-BirA*-mediated α_2A_-AR biotinylation by Rab29TN. HEK293 cells were transfected with HA-α_2A_-AR and Myc-Sar1B-BirA*, along with empty vectors or Rab29TN. **(G)** Quantitative data shown in (E) and (F). In each panel, similar results were obtained in at least 3 independent replicates. Input lanes contain 3% of the total protein used in the experiments. All quantitative data are presented as means ± SE; n = 3-4. ***p < 0.001 versus control.

We then measured the effects of Rab29 KO and inhibition on α_2A_-AR biotinylation by Sar1B- BirA*. In control cells expressing Sar1B-BirA*, α_2A_-AR was biotinylated; however, Sar1B-BirA*-mediated α_2A_-AR biotinylation was attenuated by approximately 50% in Rab29 KO cells (Figure 4E and 4G) and in cells expressing the inactive Rab29TN mutant (Figure 4F and 4G). These data suggest that similar to AT2R, α_2A_-AR targeting to COPII vesicles is regulated by Rab29.

### Rab29 localizes to the ER and COPII vesicles in an activation- and geranylgeranylation- dependent manner and directly interacts with Sec24

Rab29 is well known to localize predominantly at the Golgi, with its membrane association mediated by geranylgeranylation on two C-terminal (CT) cysteine residues (CC) (Nagai-Ito et al., 2022). To elucidate how Rab29 mediates cargo recruitment to COPII vesicles, we first determined whether it localizes to the ER and COPII vesicles. In addition to wild-type (WT) Rab29, we examined the inactive Rab29TN and Rab29DA mutants, as well as a geranylgeranylation-deficient Rab29CCSS mutant in which the CC motif was mutated to SS. Although Rab29Q67L (QL) mutant is predicted to be constitutively active and GTP-bound, it exhibited a largely diffuse cytosolic distribution, similar to the inactive TN and DA mutants (Figure 5 – figure supplement 1) (Beilina et al., 2014; Liu et al., 2018). Thus, the QL mutant was excluded from further cell-based assays in this study.

To evaluate ER localization, we performed live-cell bioluminescence resonance energy transfer (BRET) assays by co-expressing Rluc8-tagged Rab29 (or its mutants) with Venus-tagged protein tyrosine phosphatase 1B (PTP1B) as an ER marker. The BRET signal was much stronger in cells expressing WT Rab29 than in those expressing any of the Rab29 mutants (Figure 5A). To confirm the proximity between endogenous Rab29 and the ER protein C1orf27, we performed *in situ* proximity ligation assays (PLAs). The PLA signal was 10-fold greater in cells expressing GFP-C1orf27 compared with GFP-expressing control cells (Figure 5B and 5C). In addition, the Pearson’s correlation coefficient between Rab29 and C1orf27 was markedly higher than that between the Rab29CCSS mutant and C1orf27 (Figure 5 – figure supplement 1).

**Figure 5.**
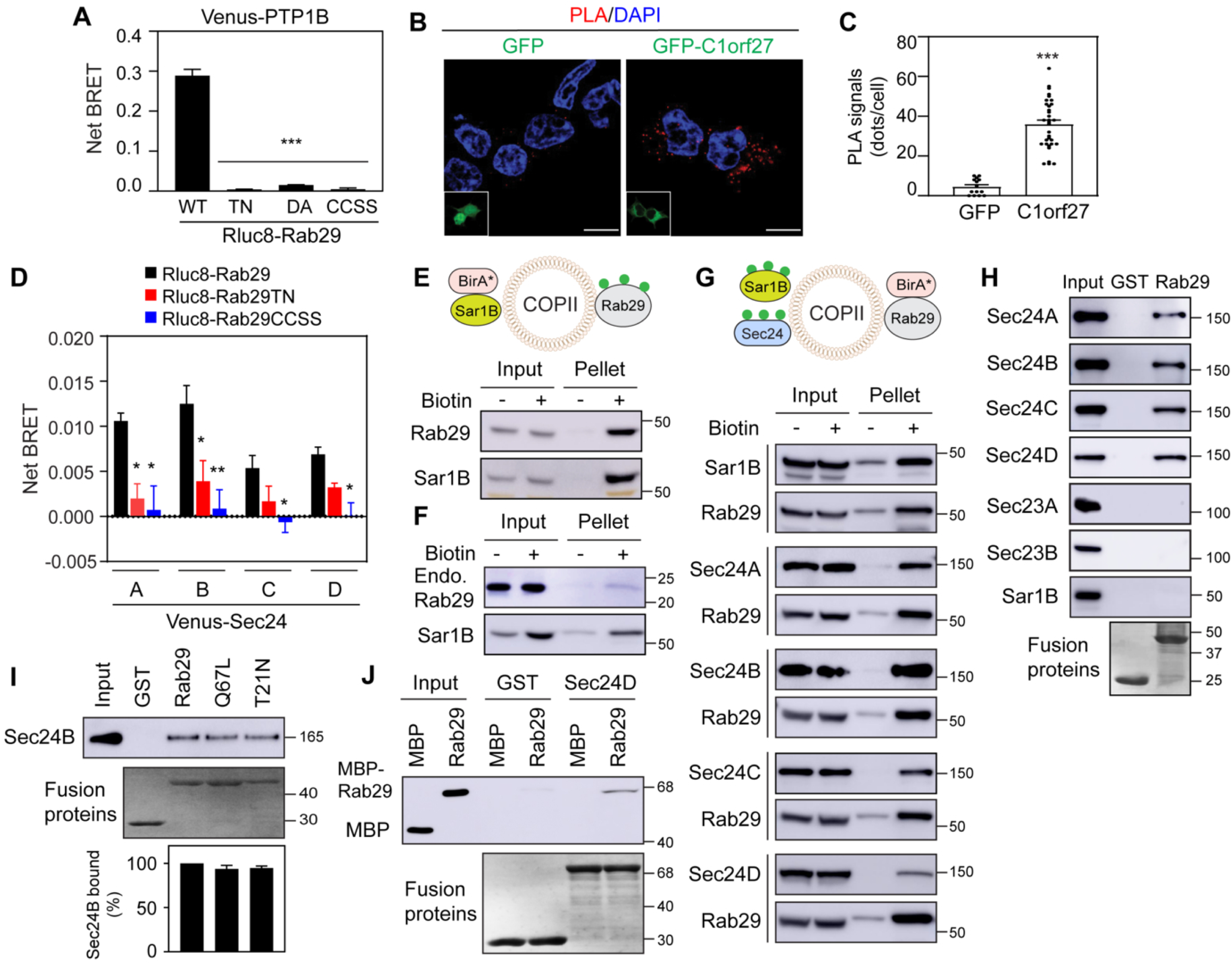
Rab29 localizes to the ER and COPII vesicles in an activation and geranylgeranylation dependent fashion and directly interacts with Sec24. **(A)** Live-cell BRET. HEK293 cells were transfected with Venus-PTP1B and Rluc8-Rab29 or its mutants. **(B)** PLA signals (red puncta) indicating proximity between endogenous Rab29 and GFP-C1orf27. HeLa cells transfected with GFP or GFP- C1orf27 were stained with antibodies against GFP and Rab29. Insets show the expression of GFP or GFP-C1orf27. **(C)** Quantitative data shown in (B). **(D)** Live-cell BRET assays evaluating proximity between Rab29 and Sec24. HEK293 cells were transfected with individual Venus-tagged Sec24 isoforms and Rluc8-tagged Rab29 or its mutants. **(E)** Rab29 biotinylation by Sar1B-BirA*. HEK293 cells were co-transfected with Myc-Sar1B-BirA* and GFP-Rab29 for 24 h and treated with biotin for 8 h. Rab29 biotinylation and total Myc-Sar1B-BirA* levels were determined using anti-GFP and anti-Myc antibodies, respectively. **(F)** Biotinylation of endogenous Rab29 by Sar1B-BirA*. **(G)** Biotinylation of Sar1B and Sec24 by Rab29-BirA*. HEK293 cells were transfected with Rab29-BirA*-HA, along with GFP-tagged Sar1B or Sec24. Biotinylation of Sar1B and Sec24 was detected using GFP antibodies, whereas Rab29-BirA*-HA expression was analyzed using anti-HA antibodies. **(H)** Rab29 interactions with COPII components in GST pulldown assays. GST-Rab29 fusion proteins were incubated with cell lysates expressing GFP-tagged Sec24, Sec23, or Sar1B. Bound proteins were detected by immunoblotting using anti-GFP antibodies. **(I)** Nucleotide state-independent interaction of Rab29 variants with Sec24. GST fusion proteins containing WT Rab29, Rab29QL, or Rab29 TN were incubated with cell lysates expressing GFP-Sec24B. The bottom panel shows the quantitative data. **(J)** Direct interaction between Rab29 and Sec24D. The Sec24D fragment (residues L^521^-N^1032^) was generated as GST fusion proteins and incubated with purified MBP or MBP-Rab29. Bound proteins were detected by immunoblotting using anti-MBP antibodies. Similar results were obtained in 3-5 individual experiments in (E)-(J). Input lanes contain 2% of the total protein used in the experiments. All quantitative data are presented as means ± SE; n = 3 independent experiments performed in triplicate in (A); n = 13-33 cells from 4 repeats in (C); n = 4 experiments performed in triplicate in (D); and n = 3 in (I). *p < 0.05 and **p < 0.01 versus Rluc8-Rab29 in (D); ***p < 0.001 versus Rluc8-Rab29 in (A) and versus GFP in (C).

We next assessed the proximity of Rab29 to COPII machinery components using both live-cell BRET and proximity-dependent biotinylation assays. In the BRET assays, Venus-tagged Rab29 (or its TN and CCSS mutants) was coexpressed with individual Rluc8-tagged Sec24 isoforms. All four Sec24 isoforms yielded substantially higher BRET signals when paired with WT Rab29 than with the Rab29 mutants (Figure 5D). In proximity biotinylation assays, both exogenously expressed (Figure 5E) and endogenous Rab29 (Figure 5F) were clearly biotinylated in cells expressing Sar1B-BirA*. Reciprocally, Sar1B and all four Sec24 isoforms were biotinylated in cells expressing Rab29-BirA* (Figure 5G). These data demonstrate that Rab29 localizes to the ER and the cytoplasmic face of COPII vesicles in an activation- and geranylgeranylation-dependent manner.

To determine whether Rab29 physically associates with COPII components, we performed GST fusion protein pulldown assays. GST-Rab29 clearly pulled down all four Sec24 isoforms, but not two Sec23 isoforms or Sar1B, whereas GST alone did not pull down any of the COPII components tested (Figure 5H). To test whether the Rab29-Sec24 interaction is dependent on the Rab29 activation status, we generated WT Rab29 along with its TN and QL mutants as GST fusion proteins and measured their binding to Sec24B. The QL mutant was included here because its membrane-association defect does not impair its binding capacity in cell-free, *in vitro* assays as suggested (Zhu et al., 2023). GST fusion proteins containing WT Rab29, Rab29QL, or Rab29TN pulled down Sec24B with similar efficiency (Figure 5I). To ascertain whether this interaction is direct, the C-terminal fragment (residues L^521^-N^1032^) of Sec24D, which contains the majority of its cargo-binding domains (Downes and Zanetti, 2025; Mancias and Goldberg, 2008), was generated as a GST fusion protein and incubated with purified maltose-binding protein (MBP)-tagged Rab29. GST-Sec24D clearly bound to MBP-Rab29, but not to MBP alone (Figure 5J). Taken together, these data demonstrate that Rab29 directly interacts with Sec24 and that this interaction is independent of its nucleotide-bound state.

### Rab29 interacts with its cargo GPCRs both directly and in an activation-dependent manner

To determine whether Rab29 interacts with cargo GPCRs at the ER, we first measured the proximity of endogenous Rab29 with individual ER-anchored RUSH GPCRs using PLAs. The PLA signals were markedly elevated in cells expressing RUSH α_2A_-AR and AT2R compared with control cells, whereas the signal in CXCR4-transfected cells remained at background levels (Figure 6A and B). We next compared the interactions of Rab29 with the intracellular domains of these three receptors. For α_2A_-AR, we tested both its CT and its large third intracellular loop (ICL3), which are known to interact with multiple proteins involved in receptor signaling and trafficking. In GST pulldown assays, the α_2A_-AR CT pulled down Rab29, whereas its ICL3 did not (Figure 6 – figure supplement 1). Consistently, the CTs of α_2A_-AR and AT2R, but not CXCR4, successfully pulled down GFP-Rab29 (top panel, Figure 6C) and endogenous Rab29 (middle panel, Figure 6C) from cell lysates. Furthermore, progressive truncation of the AT2R CT revealed that a shorter 23-residue fragment (CT_23_), but not the last 16 residues (CT_16_), retained the capacity to interact with Rab29 (Figure 6D).

**Figure 6.**
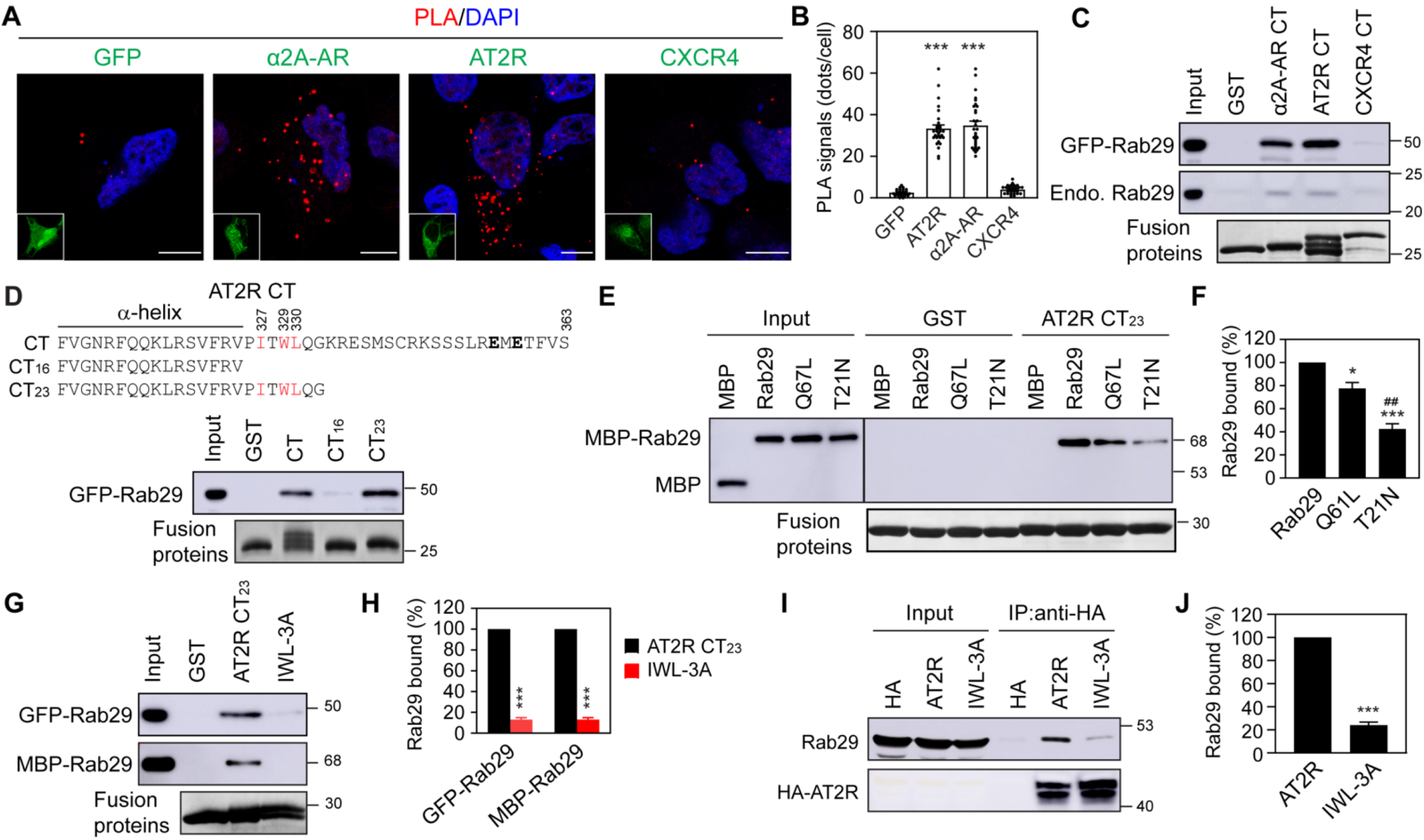
Rab29 directly and activation-dependently interacts with its cargo GPCRs. **(A)** PLA signals (red puncta) showing proximity between endogenous Rab29 and RUSH GPCRs. HEK293 cells were transfected with empty GFP vectors or GFP-tagged RUSH GPCRs and then stained with antibodies against GFP and Rab29. Insets show the expression of GFP or the GFP-tagged GPCRs. **(B)** Quantitative data shown in (A). **(C)** Interaction of the CTs of α_2A_-AR, AT2R, and CXCR4 with GFP- Rab29 (top panel) and endogenous Rab29 (middle panel) in GST pulldown assays. **(D)** Sequences of the truncated AT2R CT fragments (upper panel) and their interactions with Rab29 in GST pulldown assays (lower panel). The IxWL and ExE motifs are highlighted in red and bold, respectively. **(E)** Direct interaction of the AT2R CT_23_ fragment with MBP, MBP-Rab29, MBP-Rab29QL and MBP-Rab29TN. **(F)** Quantitative data shown in (E). **(G)** Interaction of the AT2R CT_23_ and its IWL-3A mutant with GFP-Rab29 (top panel) and purified MBP-Rab29 (middle panel). **(H)** Quantitative data shown in (G). **(I)** Co- IP of Rab29 with full-length AT2R or its IWL-3A mutant. HEK293 cells were co-transfected with GFP- Rab29 and empty HA vectors (HA), HA-AT2R, or HA-IWL-3A mutant, followed by IP using anti-HA antibodies. **(J)** Quantitative data shown in (I). Similar results were obtained in 3 individual experiments in (C). Input lanes contain 2% of the total protein used in the experiments. All quantitative data are presented as means ± SE; n = 24-30 cells from 3 experiments in (B); n = 4 in (F); n = 3 in (H) and (J). *p < 0.05 versus Rab29 in (F); ***p < 0.001 versus GFP and CXCR4 in (B), Rab29 in (F), AT2R CT_23_ in (H), and AT2R in (J); _##_ p < 0.001 versus Q67L in (F). Scale bars,10 µm.

We next determined whether the interaction between Rab29 and the GPCR domains is direct and activation-dependent using GST pulldown assays. In cell lysates, the CTs of α_2A_-AR and AT2R interacted with WT Rab29 more strongly than with the inactive TN mutant, whereas neither the CXCR4 CT nor the α_2A_-AR ICL3 interacted with either Rab29 variant (Figure 6 – figure supplement 1). To test for direct interaction, we incubated the AT2R CT_23_ fragment with purified proteins. AT2R CT_23_ pulled down purified MBP-Rab29, but not MBP alone (Figure 6E). The binding of AT2R CT_23_ to MBP-Rab29 was moderately but significantly stronger than its binding to the constitutively active MBP-Rab29QL mutant, and both interactions were substantially greater than that observed with the inactive MBP- Rab29TN mutant (Figure 6E and 6F). A similar activation-dependent binding (WT Rab29 > QL >> TN) was observed for the full-length AT2R CT (Figure 6 – figure supplement 1). Together, these data demonstrate that Rab29 directly interacts with its cargo GPCRs in an activation-dependent manner.

Because our progressive deletion data indicated that the PITWLQG sequence within the AT2R CT is critical for Rab29 binding (Figure 6D), we mutated three hydrophobic residues, I^327^, W^329^ and L^330^, to map the precise binding motif. Mutation of this IWL triplet to three alanines (IWL-3A) almost completely abolished the AT2R CT binding to both GFP-Rab29 in cell lysates and purified MBP-Rab29 in GST pulldown assays (Figure 6G and 6H). In contrast, mutating the acidic ExE motif, which was previously identified to mediate AT2R concentration into COPII vesicles and ER export (Xu et al., 2023; Zhang et al., 2011), did not alter the AT2R CT interaction with Rab29 (Figure 6 – figure supplement 1). Further, co-immunoprecipitation (co-IP) assays confirmed that the amount of Rab29 pulled down by full-length AT2R was profoundly reduced by the IWL-3A mutation (Figure 6I and 6J). These data demonstrate that AT2R interaction with Rab29 is mediated specifically by its CT IxWL motif.

We next utilized RUSH assays to determine the functional consequences of disrupting the IxWL motif on the COPII recruitment and ER-Golgi-surface transport of AT2R. Compared with its WT counterpart, the accumulation of the AT2R IWL-3A mutant within Sec24B-positive COPII vesicles was severely impaired (Figure 7A and 7B), resulting in a roughly 45% reduction in cargo intensity (Figure 7C). Consistent with this defect, the mutation clearly slowed down the ER-to-Golgi transport kinetics, with increased t_1/2_ in live-cell RUSH assays (Figure 7D-7F), as well as in fixed cells after biotin induction for 15 min (Figure 7G and 7H). Finally, RUSH-based Nano-Glo HiBiT extracellular detection assays revealed that the surface expression of the IWL-3A mutant was significantly lower compared with WT AT2R (Figure 7I). These data suggest that the direct interaction between AT2R and Rab29 governs receptor targeting to COPII vesicles and drives efficient anterograde export from the ER to the cell surface.

**Figure 7.**
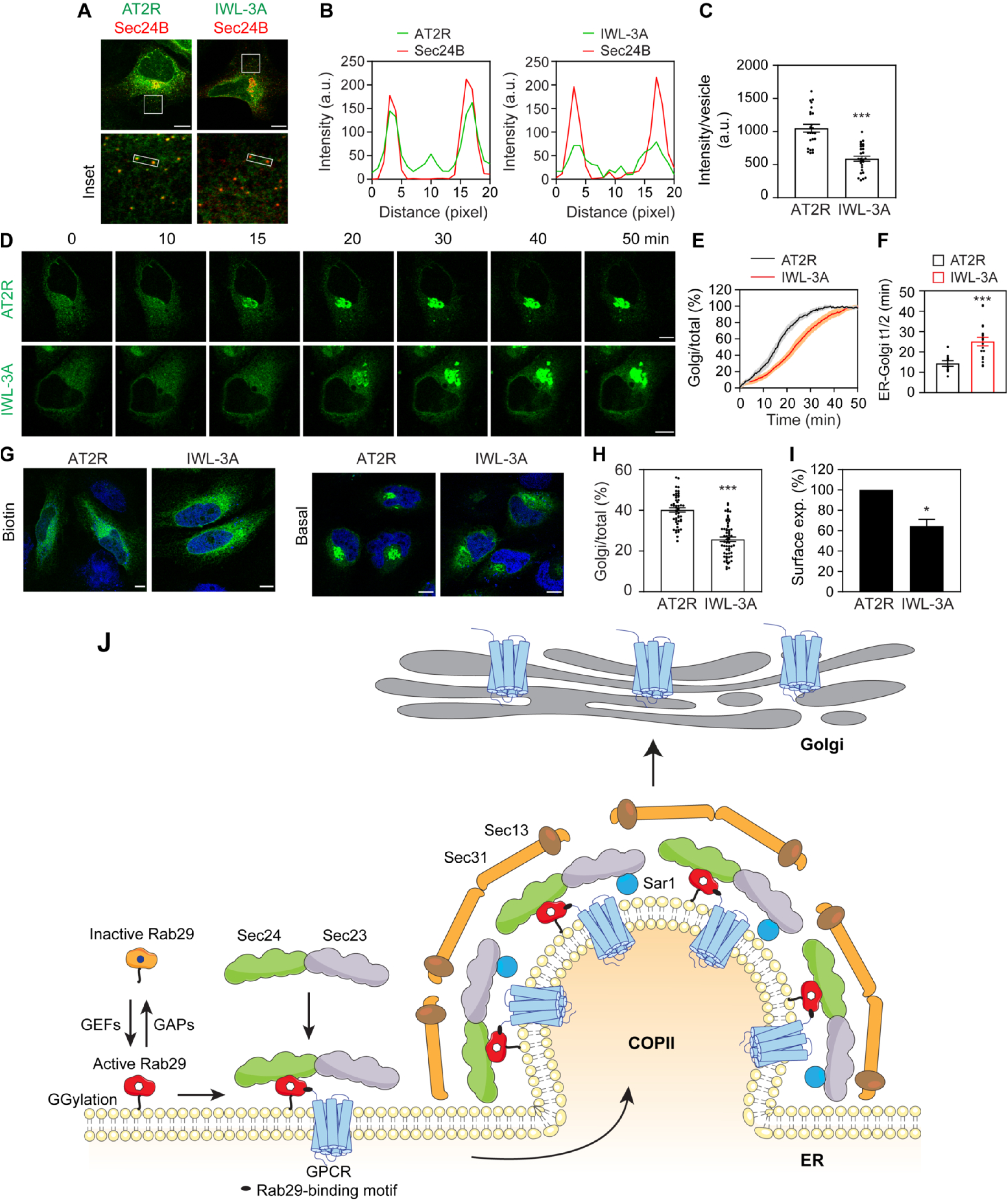
Mutation of the IxWL motif disrupts COPII concentration and ER-Golgi-surface transport of AT2R. **(A)** Representative images showing the colocalization of AT2R or its IWL-3A mutant with Sec24B. HeLa cells were transfected with GFP-tagged RUSH AT2R or its mutant, along with DsRed-Sec24B. **(B)** Analysis of AT2R and Sec24B colocalization shown in (A). **(C)** Quantitative data showing the recruitment of AT2R and its mutant to COPII vesicles. **(D)** Live-cell imaging showing the effects of mutating the IxWL motif on the export kinetics of AT2R. **(E)** The transport curves of AT2R and its mutant in HeLa cells. **(F)** t_1/2_ values of ER-to-Golgi transport for AT2R and its mutant. **(G)** ER-to- Golgi transport of AT2R and its mutant in fixed cells after biotin induction for 15 min. **(H)** Quantitative data shown in (G). **(I)** Cell surface expression of AT2R and its mutant, as quantified by Nano-Glo HiBiT assays. **(J)** A model depicting the functions and mechanism of Rab29 in facilitating the COPII recruitment of GPCRs (see text for detail). GGylation, geranylgeranylation. All quantitative data are presented as means ± SE; n = 24-27 cells in (C), 9-18 cells in (E) and (F), 49-60 cells in (H) and 4, each compiled from at least 4 individual experiments. *p < 0.05 and ***p < 0.001 versus AT2R. Scale bars,10 µm.

## Discussion

In this study, we have revealed novel functions for Rab29 in the export of newly synthesized GPCRs from the ER to the Golgi complex. These functions of Rab29 were first identified in a screening assay measuring the effects of depleting all mammalian Rabs on the ER-to-Golgi transport of nascent α_2A_-AR in MDCK cells using RUSH assays, which were further supported by Rab29 KO and the transient expression of two inactive mutants (Rab29TN and Rab29DA) to inhibit Rab29 function in HeLa and HEK293 cells. Rab29 KO and its inactive mutants markedly impeded the ER-to-Golgi transport of a specific subset of GPCRs, without affecting the transport of non-GPCR PM proteins. These transport deficits were consistently observed in both fixed and live cells. Furthermore, WT Rab29 successfully rescued these transport deficits, whereas the inactive mutants neither ameliorated nor further exacerbated the transport in Rab29 KO cells. Consistent with these trafficking defects, Rab29 inhibition reduced the steady-state cell surface expression of both exogenously transfected and endogenously expressed GPCRs and suppressed their downstream cellular responses. These functional deficits included impaired α_2A_-AR-mediated ERK1/2 activation, β_2_-AR-stimulated cAMP production, and AT2R-dependent cell proliferation inhibition. These data strongly suggest that the observed effects are likely specific. Together with previous findings (Doly et al., 2016; Petaja-Repo et al., 2000; Wei et al., 2019; Xu et al., 2024b), our data reinforce the concept that ER-Golgi transport is the rate-limiting step in GPCR maturation and targeted delivery to their functional destination. However, we cannot entirely exclude the possibility that Rab29 inhibition alters the trafficking of other signaling components, resulting in aberrant cellular responses to receptor activation. Additionally, whether Rab29 also regulates the post-Golgi transport of GPCRs or other PM proteins remains an open question.

We have identified ER-derived COPII vesicles as the site of action where Rab29 enhances the capture of nascent GPCRs. Direct evidence for Rab29 function at the COPII level includes a marked reduction in the concentration and fluorescence recovery dynamics of AT2R within COPII vesicles in Rab29 KO and inactive mutant-expressing cells, as measured by imaging and live cell FRAP. Additionally, Rab29 inhibition significantly attenuated Sar1B-BirA*-mediated biotinylation of α_2A_-AR, further suggesting a role for Rab29 in recruiting α_2A_-AR to COPII vesicles. Rab29 function at COPII vesicles is supported by its localization at both the ER and COPII vesicles, as revealed by live-cell BRET, PLAs, colocalization, and proximity biotinylation assays. These experiments demonstrated the close spatial proximity of Rab29 to ER resident proteins (C1orf27 and PTP1B) and COPII components (Sec24A-D and Sar1B). Specifically, the live-cell BRET signals for individual Sec24 isoforms were substantially higher in cells expressing WT Rab29 compared with those expressing inactive or geranylgeranylation-deficient mutants. Furthermore, Sar1B-BirA* successfully biotinylated Rab29, and Rab29-BirA* reciprocally biotinylated all four Sec24 isoforms and Sar1B. Together, these data demonstrate that Rab29 localizes to the cytoplasmic face of COPII vesicles in an activation- and geranylgeranylation-dependent manner, aligning with the prior proteomic identification of Rab29 in streptavidin pulldowns of Sar1-BirA*-biotinylated proteins (Emmer et al., 2018).

Our study has elucidated the molecular mechanism underlying Rab29 function in GPCR export, which is mediated through direct interactions with Sec24 and cargo GPCRs (Figure 7J). Consistent with their proximity measured in live-cell BRET and biotinylation assays, Rab29 physically associated with all four Sec24 isoforms in GST pulldown assays. By contrast, Rab29 did not interact with Sar1B (despite their mutual biotinylation), Sec23A, or Sec23B, confirming the specificity of the Rab29-Sec24 interaction. Furthermore, because the GST fusion protein containing the C-terminal half of Sec24D successfully pulled down purified MBP-Rab29, the interaction between Sec24 and Rab29 is likely direct. Although WT Rab29, its active QL mutant, and its inactive TN mutant bound Sec24B with similar efficiency, the COPII localization of Rab29 in intact cells strictly relied on its activation state. This implies that only the active form of Rab29 can interact with Sec24 at COPII vesicles *in vivo*. These findings align with established paradigm that while Sec23 interacts with activated Sar1 to promote COPII vesicle biogenesis, Sec24 interacts with cargoes or cargo receptors to enhance cargo capture by COPII vesicles.

In addition to interacting with Sec24, Rab29 also interacts with cargo GPCRs. Importantly, these interactions are selective, direct, and dependent on the activation of Rab29. Our data revealed robust PLA signals for endogenous Rab29 in cells expressing the Rab29-regulated receptors AT2R and α_2A_- AR, but not in cells expressing the Rab29-independent receptor CXCR4. Furthermore, Rab29 interacted with the CTs of α_2A_-AR and AT2R, but not the CXCR4 CT, indicating that Rab29 selectively interacts with the specific GPCRs it regulates. The direct and activation-dependent nature of these interactions was demonstrated in GST pulldown assays, where the CTs of α_2A_-AR and AT2R pulled down GFP-Rab29 and purified MBP-Rab29 much more efficiently than the inactive TN mutant. This activation-dependent binding is further supported by the fact that the Rab29QL mutant, which possesses a lower GTP-binding affinity (Beilina et al., 2014), exhibited reduced binding to the AT2R CT compared with its WT counterpart. Additionally, we identified a specific hydrophobic IxWL motif that mediates AT2R interaction with Rab29, as validated in GST pulldown and co-IP assays; notably, this motif is absent in the intracellular domains of the other five Rab29-regulated GPCRs. Mutating this motif significantly impaired COPII concentration, ER-to-Golgi transport, and cell surface expression of AT2R, highlighting a crucial role for the Rab29 interaction in controlling AT2R export from the ER via COPII vesicles. However, the specific motif mediating the α_2A_-AR-Rab29 interaction, and whether other Rab29-regulated GPCRs interact with Rab29, remain to be determined.

Based on the data presented in this study, we propose a model wherein activated Rab29 localizes to the ER membrane and directly interacts with both Sec24 and cargo GPCRs, thereby enhancing GPCR recruitment into COPII vesicles for subsequent delivery to the Golgi complex (Figure 7J). Thus, the Rab29 GTPase functions as a selective cargo receptor for the ER export of specific nascent GPCRs. There are several important distinctions regarding Rab29 acting as a cargo receptor. First, previously identified cargo receptors are ER-localized transmembrane proteins, such as ERGIC53 (also known as lectin mannose-binding protein 1) (Appenzeller et al., 1999), transmembrane Emp24 domain-containing (TMED) proteins (Barlowe and Helenius, 2016), surfeit locus protein 4 (Emmer et al., 2018; Huang et al., 2021; Tang et al., 2022; Wang et al., 2021b), Wolfram syndrome 1 (Wang et al., 2021a), Cornichon proteins (Zhang and Schekman, 2016), and CLN8 (di Ronza et al., 2018). The cytosolic protein 14-3-3 can also act as a cargo adaptor to facilitate ER export (Bajaj Pahuja et al., 2015), These proteins lack enzymatic activity and are not regulated by nucleotide activation. In contrast, Rab29 is a small GTPase whose function strictly depends on its activation state and ER membrane association. However, specific GEFs and GAPs for Rab29 are unknown, and the mechanisms governing its activation and inactivation *in vivo* remain poorly defined. Second, while several Rab GTPases interact with cargo molecules (Carroll et al., 2001), including GPCRs (Li et al., 2012; Li et al., 2017), Rab1b is the only other Rab GTPase shown to interact with COPII components (Slavin et al., 2011). Rab1 is the founding member of the Rab family and is highly conserved across eukaryotes (Carroll et al., 2001; Mizuno-Yamasaki et al., 2012; Stenmark, 2009). It mediates the ER-to-Golgi transport of nearly all proteins by regulating multiple sequential steps, including COPII vesicle formation, motility, tethering, and fusion (Allan et al., 2000; Slavin et al., 2011). In marked contrast, Rab29 inhibition neither altered COPII vesicle formation (as measured by vesicle number and Sec24B intensity) nor globally disrupted the ER export of all cargo molecules. This indicates that Rab29 does not affect the general machinery essential for the transport of diverse cargoes. Third, while studies over the past few decades have identified numerous proteins that interact with specific GPCRs to enhance their biosynthetic transport from the ER (Zhang and Wu, 2019), including Cornichon proteins (Sauvageau et al., 2014) and TMED proteins (Di Minin et al., 2022), which are proposed to function as cargo receptors, none have been clearly proven to act on GPCRs specifically at the level of COPII vesicles. As such, to the best of our knowledge, Rab29 represents the first GTPase and the only GPCR-interacting protein demonstrated to serve as a cargo transport receptor that directly facilitates GPCR recruitment into COPII vesicles.

It is increasingly apparent that the anterograde trafficking of nascent GPCRs from the ER to their functional destinations, as well as their segregation from one another and from other PM proteins, is a highly coordinated process regulated by distinct factors at each transport stage. The data presented here, alongside our previous studies demonstrating the role of the ufmylation system in the COPII recruitment of specific GPCRs (excluding AT2R) (Xu et al., 2026; Xu et al., 2024a), indicate that distinct GPCRs utilize different mechanisms for COPII vesicle targeting. These distinct pathways may provide crucial checkpoints for the selective control of receptor function. It is worth noting that, compared to other well-studied GPCRs like AT1R, the physiological functions of AT2R remain largely unexplored. However, a growing body of evidence indicates that AT2R is an attractive therapeutic target for hypertension, obesity, neuropathic pain, lung fibrosis, and glioblastoma (Steckelings et al., 2022; Zhang et al., 2021). Our finding that Rab29 inhibition impairs AT2R-mediated antiproliferation suggests that targeting receptor recruitment to COPII vesicles represents a viable approach to modulating AT2R functions. Furthermore, because numerous GPCRs, including adrenergic and Ang II receptors regulated by Rab29, are implicated in the pathogenesis of PD and serve as clinical targets for its treatment (Jones-Tabah, 2023), a thorough elucidation of the functional relationship between Rab29 and neuronal GPCRs may provide a foundation for therapeutic innovation.

In summary, our study has identified a novel function for the Rab family in membrane trafficking. Rab29 represents the first GTPase that functions as a selective cargo receptor for newly synthesized GPCRs. These findings provide important insights into COPII-mediated protein transport and the regulation of GPCR biosynthetic maturation

## Materials and methods

### KEY RESOURCES TABLE

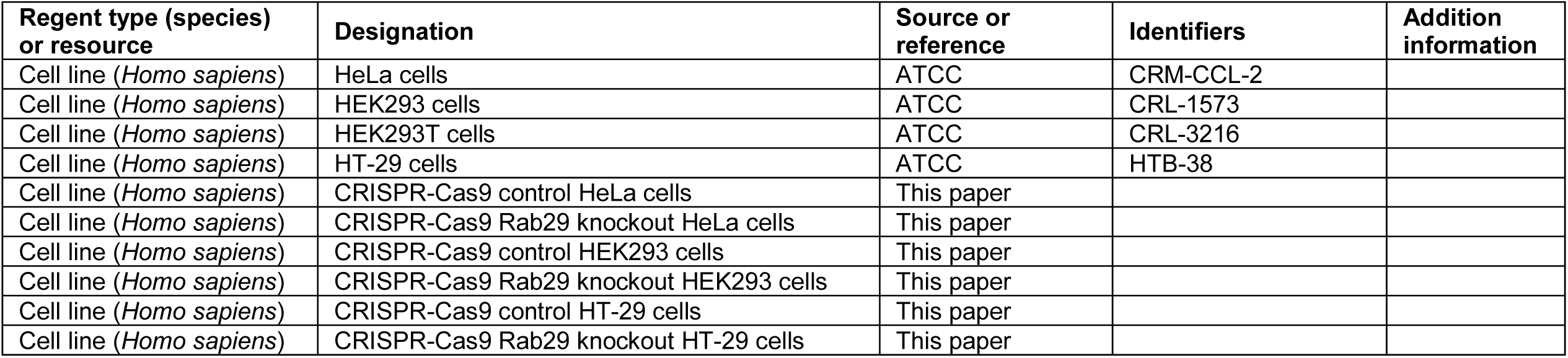
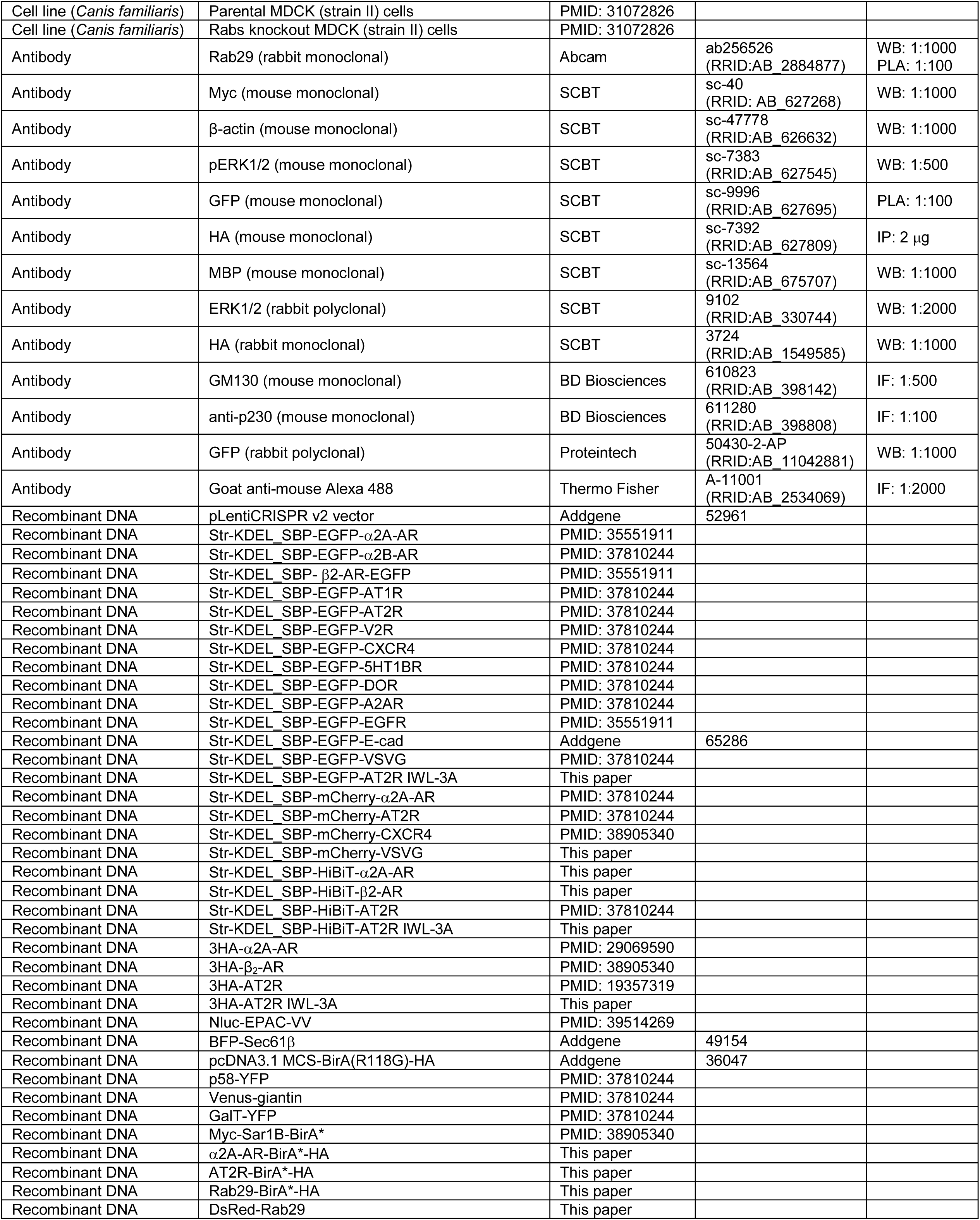
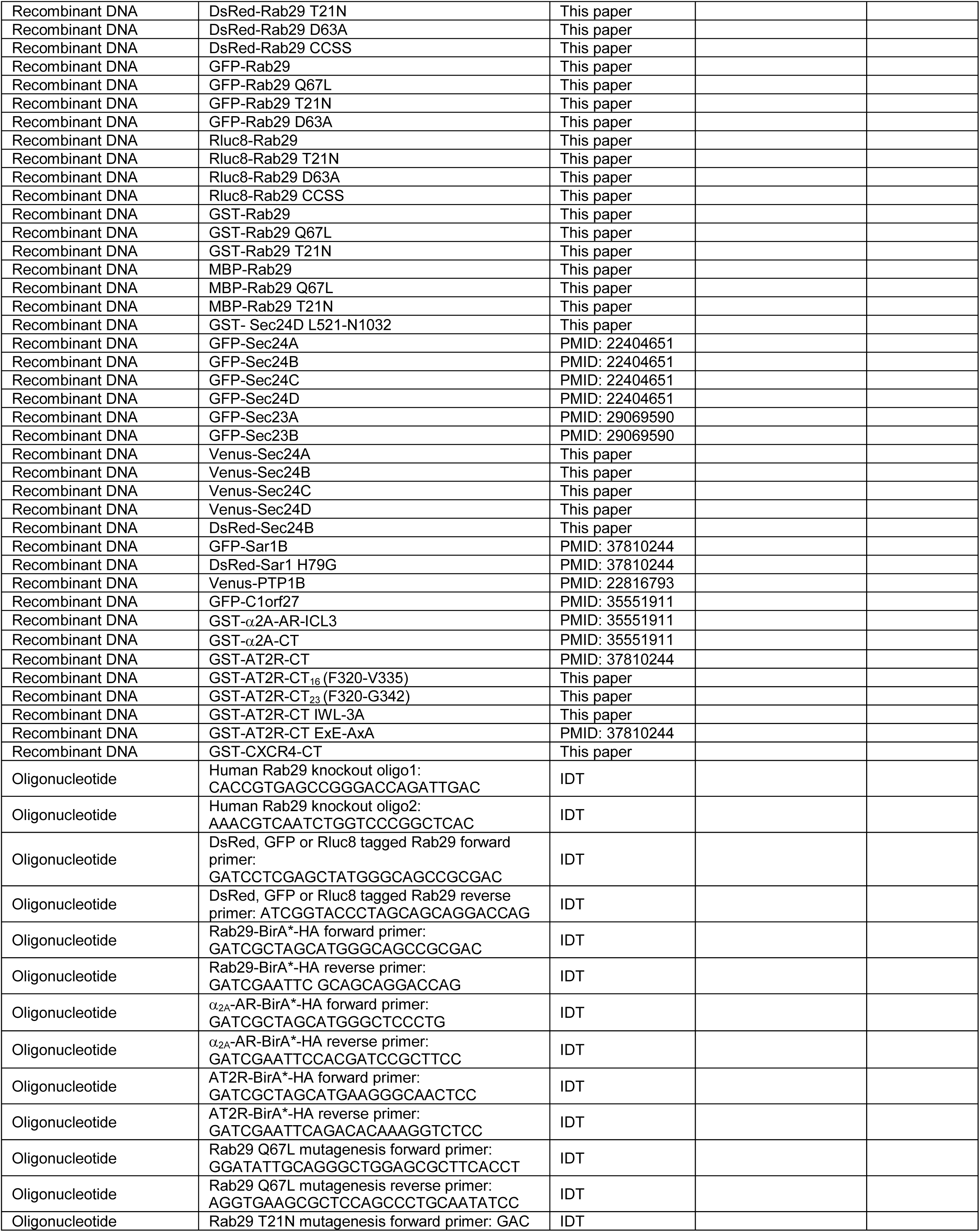
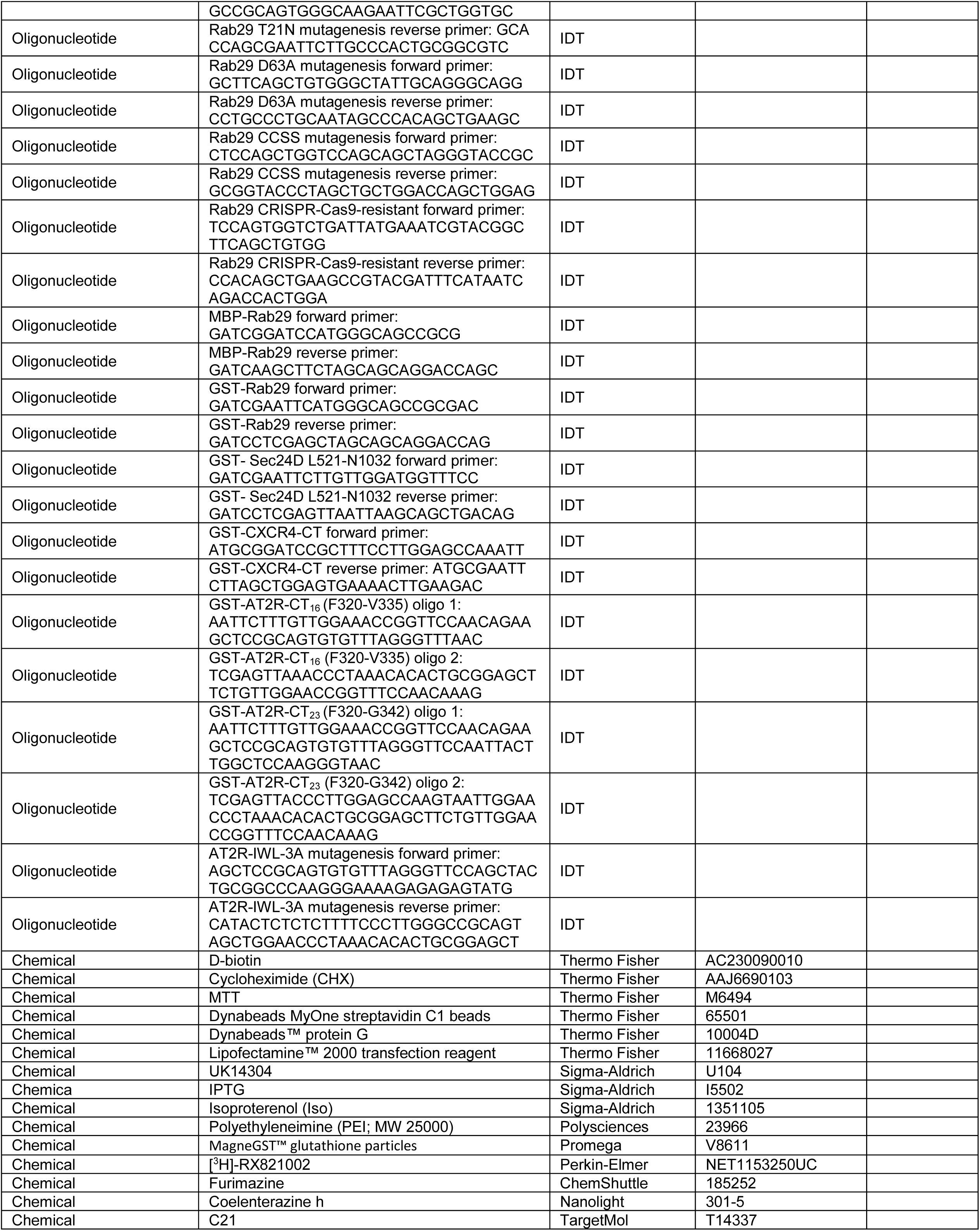
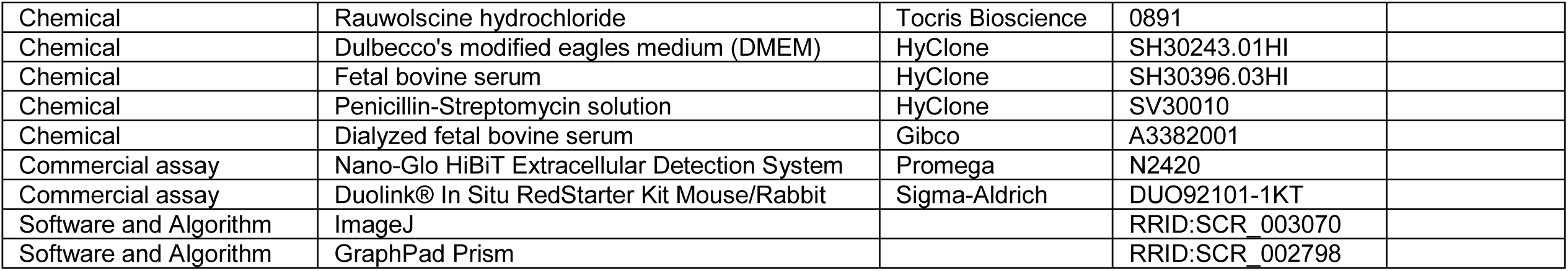

### Plasmids and construction

RUSH plasmids expressing α_2A_-AR, α_2B_-AR, β_2_-AR, AT1R, AT2R, V2R, CXCR4, 5HT1R, DOR, A2AR, EGFR, E-cadherin, and VSVG tagged with GFP, mCherry, or HiBiT, plasmids expressing 3HA-tagged α_2A_-AR and AT2R, GFP- or Venus-tagged Sec24A, Sec24B, Sec24C, and Sec24D, GFP-tagged Sec23A, Sec23B, Sar1B, and C1orf27, DsRed-Sar1BH79G, and Myc- Sar1B-BirA* were generated as described previously (Xu et al., 2024a; Xu et al., 2023). Rab29 in pDsRed-Monomer-C1, pEGFP-C1, and Rluc8-C1 vectors was generated using the primers and then subcloned into the Xhol and Kpnl sites. BirA*-HA-tagged Rab29 and α_2A_-AR were generated using the primers and subcloned into pcDNA3.1 MCS-BirA(R118G)-HA vectors (Addgene Cat #36047) at the Nhel and EcoRl sites. MBP-tagged Rab29 was generated using the primers and subcloned into the pMAL-c5X vector at the BamHI and HindIII sites. GST fusion protein constructs edcoding Rab29, the C- terminal fragment (residues L521-N1032) of Sec24D, and the CXCR4 CT were generated in pGEX-4T- 1 vectors using the primers. To generate GST-tagged constructs of different lengths of the AT2R CT and its mutants, double-stranded DNA fragments were synthesized and subcloned into pGEX-4T-1 vectors. All mutagenesis was carried out using the QuikChange site-directed mutagenesis kit (Agilent) and the following primers. All constructs used in the present study were verified by nucleotide sequence analysis.

### Cell culture and transfection

HeLa, HEK293, HEK293T, HT-29, and MDCK (strain II) cells (Homma et al., 2019) were cultured in Dulbecco’s modified Eagle’s medium (DMEM) supplemented with 10% fetal bovine serum (FBS), 100 units/ml penicillin, and 100 μg/ml streptomycin. For proximity-dependent biotinylation assays, cells were cultured in DMEM containing 10% dialyzed FBS. Transient transfection of cells was performed using PEI.

### Generation of Rab29 KO cell lines using CRISPR-Cas9 genome editing

MDCK Rab KO cells were generated as described (Homma et al., 2019). To generate Rab29 KO HeLa and HEK293 cells, the sgRNA sequence 5’-GGTCTGACTACGAGATAGTG-3’ was cloned into the pLentiCRISPR v2 vector (Addgene, plasmid #52961). Lentiviral particles expressing sgRNA and Cas9 were produced via standard transfection of HEK293T cells. Following viral infection, stable cell clones were isolated using puromycin selection and limiting dilution. Successful Rab29 KO was determined by immunoblotting.

### Fluorescence microscopy

All images were captured using LAS X software with a 63x objective on a Leica Stellaris 5 confocal microscope equipped with an Okolab UNO stage-top incubator. To monitor live-cell cargo transport, time-lapse images were acquired at 30-sec intervals as described previously (Xu et al., 2024a). To evaluate protein transport in fixed cells, the cells were fixed in 4% paraformaldehyde for 15 min prior to imaging. For subcellular distribution analysis of endogenous Rab29, GM130, or p230, HeLa cells were fixed with 4% paraformaldehyde for 15 min, permeabilized with 0.1% saponin for 20 min, and blocked with 0.24% normal donkey serum for 1 h. The cells were then incubated with primary antibodies overnight at 4 °C followed by Alexa Flour-conjugated secondary antibodies for 1 h at room temperature.

### RUSH transport assays

RUSH assays were essentially carried out as described previously (Boncompain et al., 2012; Xu et al., 2024a; Xu et al., 2024b; Xu and Wu, 2022). Briefly, cells were transfected with RUSH plasmids for 20 h and then incubated with 40 µM biotin and 400 µg/ml CHX for the durations indicated in the figure legends. Cargo expression in the Golgi and the whole cell was quantified by measuring the fluorescence intensities using NIH ImageJ software. The ER-to-Golgi transport efficient was calculated as the Golgi-to-total expression ratio. For live-cell RUSH assays, the Golgi-to-total ratio at each time point was subtracted from the baseline ratio at time 0 and normalized to the maximum ratio observed in control cells, which was defined as 100%.

### Nano-Glo HiBiT extracellular detection assays

The cell surface expression of GPCRs was quantified using the Nano-Glo HiBiT Extracellular Detection System (Promega) as described previously *(Xu et al., 2023).* Briefly, HEK293 cells cultured in 6-well plates were transfected with individual HiBiT- tagged RUSH GPCRs (500 ng/well), either alone or co-transfected with Rab29 mutants (1,500 ng/well) for 6 h. The cells were then split into 12-well plates and cultured for additional 20 h. Following a 60-min treatment with biotin at 37°C, the cells were washed twice with 1 ml of Dulbecco’s phosphate-buffered saline (DPBS) and suspended in 350 µl of DPBS. A 100-µl aliquot of cell suspension was transferred into a white 96-well plate and incubated with 50 µl of LgBiT protein (1:1,000 dilution) for 10 min at 37°C. Finally, 50 µl of furimazine substrate (1:1,000 dilution) was added to each well and luminescence was measured in a Mithras LB940 photon-counting plate reader (Berthold Technologies).

### Radioligand binding

The cell surface expression of endogenous α_2A_-AR in HT29 control and Rab29KO cells were quantified by radioligand binding using membrane preparations as described previously (Duvernay et al., 2004). Briefly, membrane suspensions containing 150 µg of protein were incubated with 2 nM [^3^H]-RX-821002 for 1 h at room temperature. Non-specific binding was determined in the presence of 20 µM rauwolscine. The reaction was terminated by vacuum filtration, and radioactivity was measured via liquid scintillation spectrometry using an AccuFLEX LSC-8000 counter (Hitachi Aloka).

### Measurement of ERK1/2 activation

α_2A_-AR-mediated ERK1/2 activation was determined by measuring phosphorylation via immunoblotting as described previously (Xu et al., 2024a). HEK293 cells cultured on 12-well plates were transfected with α_2A_-AR, either alone or together with Rab29TN for 24 h. Cells were serum-starved for 12 h and subsequently stimulated with 1 µM UK14304 for 5 min. Stimulation was terminated by adding 100 µl of SDS gel loading buffer.

### cAMP measurement

β_2_-AR-stimulated cAMP production in HEK293 cells was measured via live-cell BRET as described previously (Jang et al., 2024). HEK293 cells seeded in 6-well plates were transfected with the cAMP sensor Nluc-EPAC-VV (10 ng/well) and β_2_-AR (50 ng/well) for 20 h. Cells were then split onto white 96-well plates and incubated with 5 µM furimazine substrate. BRET signals were measured before and after the addition of 10 µM ISO using a LUMIstar Omega plate reader (BMG Labtech). BRET kinetics were calculated using MARS Data Analysis software and the data at each time point were normalized to the baseline BRET signal at time 0.

### Cell proliferation

HT29 cells were seeded in 96-well plates at a density of 500 cells/well. The cells were transfected with AT2R for 6 h and subsequently treated with C21 at various concentrations (0.1, 1 or 10 µM) for 14 days. The cell culture medium containing C21 was replenished every other day. Cell proliferation was quantified using standard MTT assays.

### FRAP assays

Control and Rab29 KO HeLa cells were cultured in 3.5-cm glass-bottom dishes and transfected with GFP-tagged RUSH GPCR plasmids (500 ng) for 20 h. Photobleaching of GFP was then performed using 488-nm laser at full power. Post-bleach images were captured at 3-sec intervals for 2 min using less than 2% laser intensity.

### Proximity-dependent biotinylation and streptavidin pulldown assays

To study the proximity of α_2A_-AR to COPII components or Rab29, HEK293 cells were seeded in 6-cm dishes and transfected with α_2A_-AR-BirA*-HA plus GFP-tagged Sec24 or Rab29 (2 µg of each plasmid). After 6 h, cells were treated with 15 µM biotin for 24 h. To study the proximity of Rab29 to COPll components, cells were transfected with Myc-Sar1B-BirA*and GFP-Rab29, or with Rab29-BirA*-HA and GFP-tagged Sar1B or Sec24 (2 µg of each plasmid) for 24 h, and subsequently treated with 15 µM biotin and 400 µg/ml CHX for 8 h. To determine the effect of Rab29TN on the COPII-mediated transport of α_2A_-AR, cells were transfected with Myc-Sar1B-BirA* and HA-α_2A_-AR, along with either empty vectors or Rab29TN for 6 h, and then treated with biotin for 24 h. Following biotinylation, cells were harvested and lysed with 300 µl of radioimmunoprecipitation assay (RIPA) buffer [50 mM Tris-HCl (pH 7.4), 150 mM NaCl, 0.25% deoxycholic acid, 1% NP-40, and 1 mM EDTA] containing 1% protease inhibitors. After centrifugation, the supernatants were incubated with 20 µl of Dynabeads MyOne Streptavidin C1 beads overnight at 4°C. The beads were washed three times with RIPA buffer, and the biotinylated proteins were detected by immunoblotting as described previously (Xu et al., 2024a).

### BRET assays

Live-cell BRET assays were used to detect protein localization in HEK293 cells as described previously (Xu et al., 2024a). To study Rab29 localization at the ER, cells cultured in 6-well plates were transfected with Rluc8-tagged Rab29 or its mutants (250 ng/well) and Venus-tagged PTP1B (750 ng/well) for 24 h. To evaluate Rab29 localization at COPII vesicles, cells were transfected with Rluc8-tagged Rab29 (250 ng/well) and individual Venus-tagged Sec24 isoforms (750 ng/well). The cells were harvested and split into black 96-well plates. Following the addition of coelenterazine h at a final concentration of 2 µM, luminescence was measured using a Mithras LB940 photon-counting plate reader. BRET signals were calculated by dividing the emission intensity at 520-545 nm by the emission intensity at 475-595 nm.

### PLAs

PLAs were performed by using the Duolink *in situ* Red Starter Mouse/Rabbit Kit (Sigma-Aldrich). To measure Rab29 localization at the ER, HeLa cells grown on coverslips in 12-well plates were transfected with 500 ng of either GFP or GFP-C1orf27 as an ER marker for 24 h. To study potential interactions between Rab29 and ER-localized GPCRs, cells were transfected with GFP-tagged RUSH GPCR plasmids (500 ng). After fixation with 4% paraformaldehyde and permeabilization with 0.1% saponin, cells were blocked with Duolink blocking solution for 1 h and incubated with mouse anti-GFP antibodies (1:200 dilution) and rabbit anti-Rab29 antibodies (1:100 dilution) overnight at 4°C. The cells were then incubated with secondary antibodies conjugated with plus and minus Duolink II PLA probes for 1 h, followed by ligation solution containing ligase for 30 min, and amplification solution containing polymerase for 2 h at 37°C. PLA puncta were counted using Image J software, and a total of 50 randomly selected cells were quantified per group

### GST fusion protein pulldown assays

GST fusion protein pulldown assays were carried out using the MagneGST pull-down system as described previously (Xu et al., 2024a). Briefly, GST fusion proteins were incubated with HEK293 cell homogenates expressing GFP-tagged Rab29, Sec24, Sec23, or Sar1B in a total volume of 400 µl binding buffer [20 mM Tris-HCl (pH 7.4), 140 mM NaCl, 1% Nonidet P-40 and 10% glycerol] overnight at 4°C. To study the interaction of endogenous Rab29, GST fusion proteins were incubated with cell lysates prepared from untransfected cells. To evaluate direct protein- protein interaction, MBP-tagged Rab29 and MBP alone were expressed in BL21, purified using amylose resin (New England Biolabs), and eluted with maltose. Purified proteins (0.5 µg per reaction) were incubated in binding buffer containing 20 mM Tris-HCl (pH 7.4), 140 mM NaCl and 4% Nonidet P- 40 overnight at 4°C. Glutathione beads were collected and washed three times with binding buffer, and the bound proteins were detected by immunoblotting.

### Co-IP

HEK293 cells were transfected with GFP-Rab29, along with empty vectors, HA-AT2R, or HA-IWL-3A, for 24 h. Cells were lysed with 500 µl of lysis buffer [50 mM Tris-HCl (pH 7.4), 150 mM NaCl, 1% Nonidet P-40, 0.5% sodium deoxycholate, 0.1% SDS and 1% protease inhibitor cocktail]. After centrifugation, the supernatants were incubated with 2 µg of anti-HA antibodies overnight at 4 ℃, followed by incubation with 20 µl of Dynabeads Protein G (Invitrogen) for 1 h at 4 ℃. Immunoprecipitated Rab29 and AT2R were detected by immunoblotting using anti-Rab29 and anti-HA antibodies, respectively.

### Statistical analysis

Statistical differences among more than two groups were analyzed using one-way analysis of variance (ANOVA) followed by Tukey’s multiple comparisons post-hoc test. For comparisons. Between two groups, an unpaired Student’s t-test was performed. All data are expressed as means ± SE or means ± SD. Significance levels are defined as \**p* < 0.05, \*\**p* < 0.01, and \*\*\**p* < 0.001.

## Acknowledgements

This work was supported by the National Institutes of Health grant R35GM136397 (to G.W.).

## Author contributions

X.X. and G.W. conceived and designed the experiments. X.X., W.H., N.A.L and G.W. performed the experiments. X.X., W.H. and G.W. analyzed the results. N.A.L and M.F. provided materials. X.X., N.A.L and G.W wrote the manuscript.

---

**Figure 1 – figure supplement 1.**
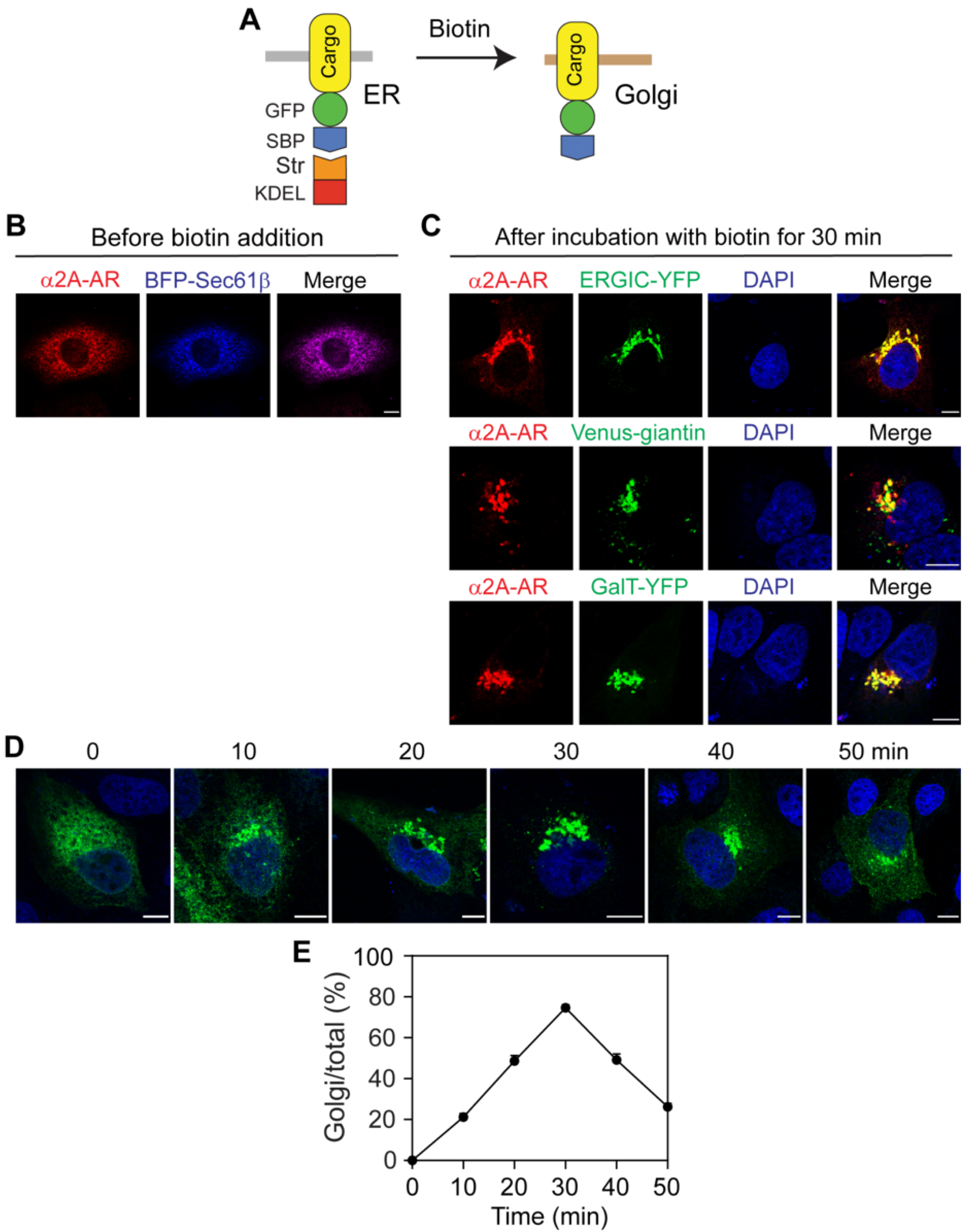
Characterization of the RUSH system to α_2A_-AR export in MDCK cells. **(A)** The RUSH system. **(B)** Colocalization of α_2A_-AR with Sec61β before biotin induction. **(C)** Colocalization of α_2A_-AR with ERGIC53, giantin and GalT after 30 min of biotin induction. Similar results were obtained in 3 separate experiments in (B) and (C). **(D)** Representative images showing the transport kinetics of RUSH α_2A_-AR in MDCK cells. **(E)** Quantitative data shown in (D). Quantitative data are presented as means ± SE; n = 20-23 cells in 3 individual experiments. Scale bars,10 µm.

**Figure 1 – figure supplement 2.**
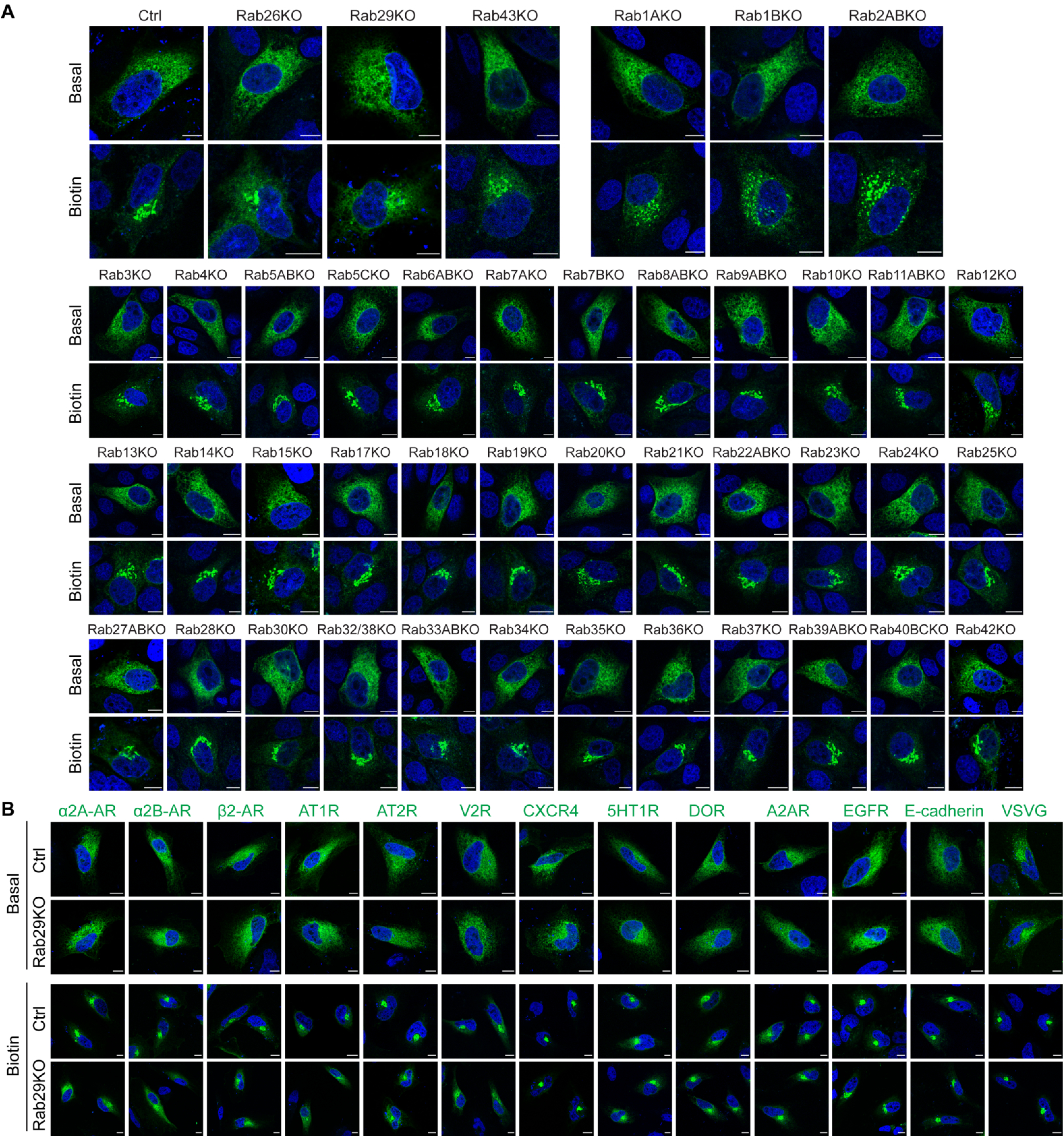
Effects of Rab KO on the ER-to-Golgi transport of GPCRs and non-GPCR proteins in MDCK and HeLa cells. **(A)** Representative images showing the effects of Rab KO on the ER-to-Golgi transport of α_2A_-AR in MDCK cells. Control and Rab KO cells were transfected with RUSH α_2A_-AR and then incubated with biotin for 15 min. **(B)** Representative images showing the effects of Rab29 KO on the ER-to-Golgi transport of different PM proteins in HeLa cells. Cells were transfected with individual RUSH cargoes and then incubated with biotin for 15 min. Scale bars,10 µm.

**Figure 1 – figure supplement 3.**
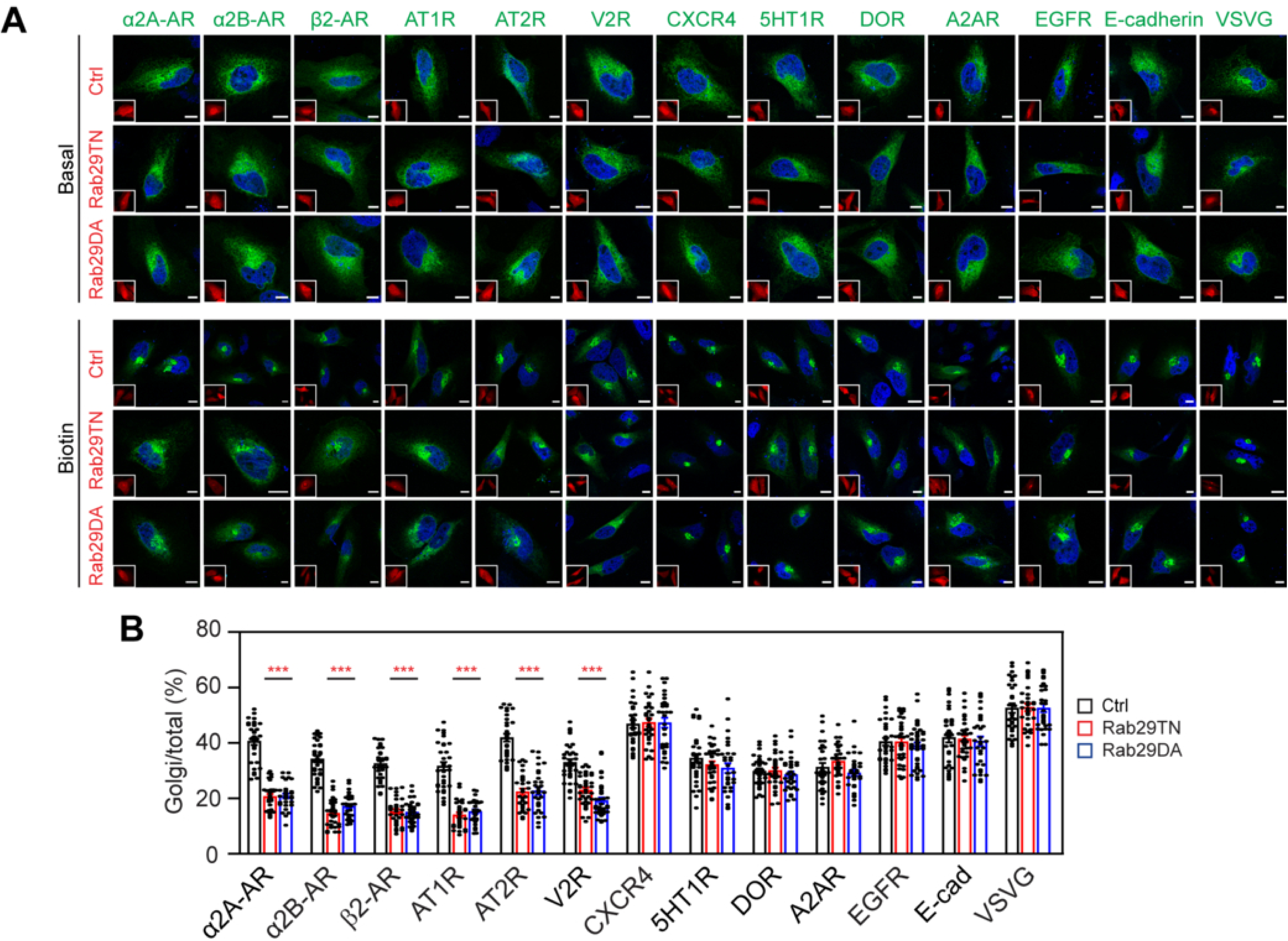
Inactive Rab29 mutants inhibit the ER-to-Golgi transport of selective GPCRs in HeLa cells. **(A)** Representative images showing the ER-to-Golgi transport of PM proteins in RUSH assays. HeLa cells were transfected with individual RUSH cargoes plus control DsRed or DsRed-Rab29 TN and DA mutants. Insets show the expression of DsRed or DsRed-Rab29 mutants. **(B)** Quantitative data shown in (A). The quantitative data are means ± SE; n = 29-41 cells from at least 4 experiments. ***p < 0.001 versus control. Scale bars,10 µm.

**Figure 1 – figure supplement 4.**
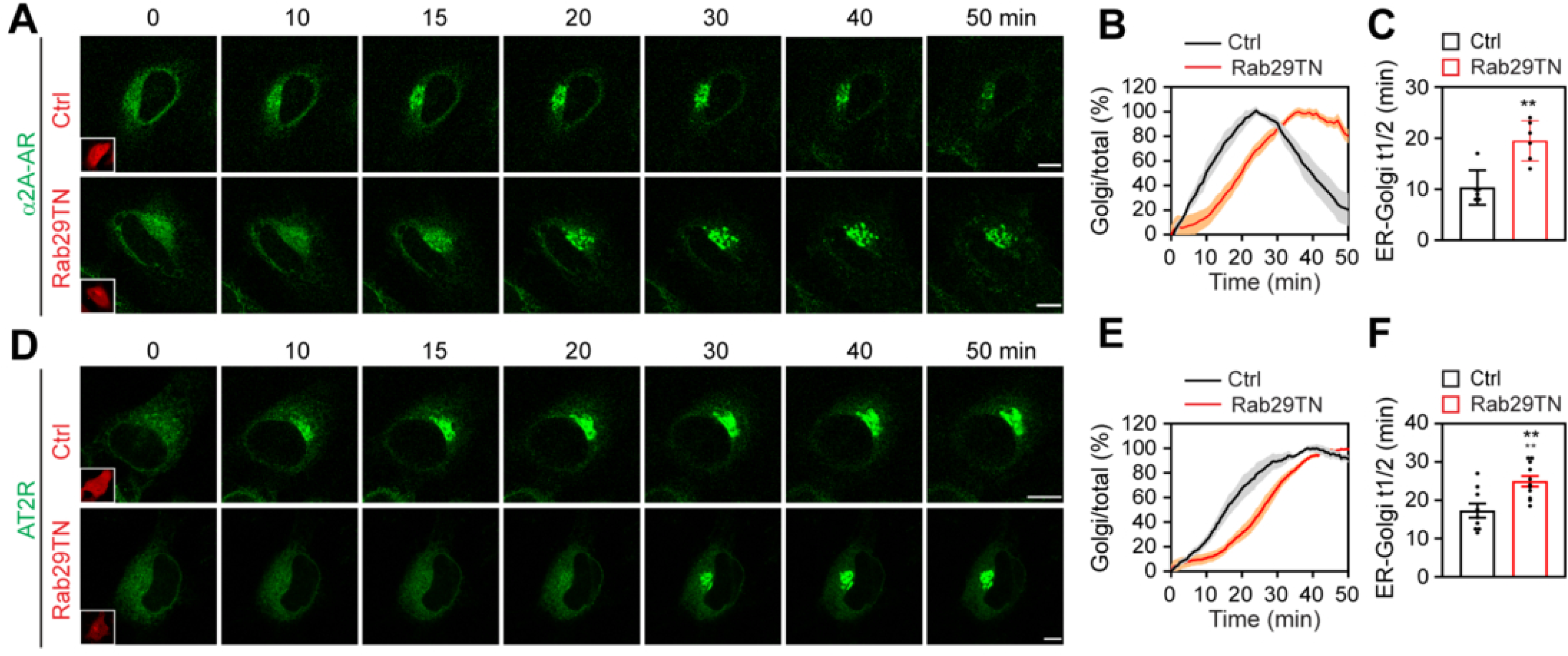
Effects of inactive Rab29 mutants on the ER export kinetics of α_2A_-AR and AT2R in live cells. **(A)** and **(D)** Live-cell imaging showing the effect of Rab29TN on the export of α_2A_-AR (A) and AT2R (D). **(B)** and **(E)** Transport curves of α_2A_-AR (B) and AT2R (E) in control and Rab29TN-expressing cells. (C) and (F) t1/2 values of ER-to-Golgi transport of α_2A_-AR (C) and AT2R (F) in control and Rab29TN-expressing cells. The quantitative data are means ± SE; n = 6 cells in (B) and (C), and 10 cells in (E) and (F), each from at least 3 experiments. **p < 0.01 versus control. Scale bars,10 µm.

**Figure 1 – figure supplement 5.**
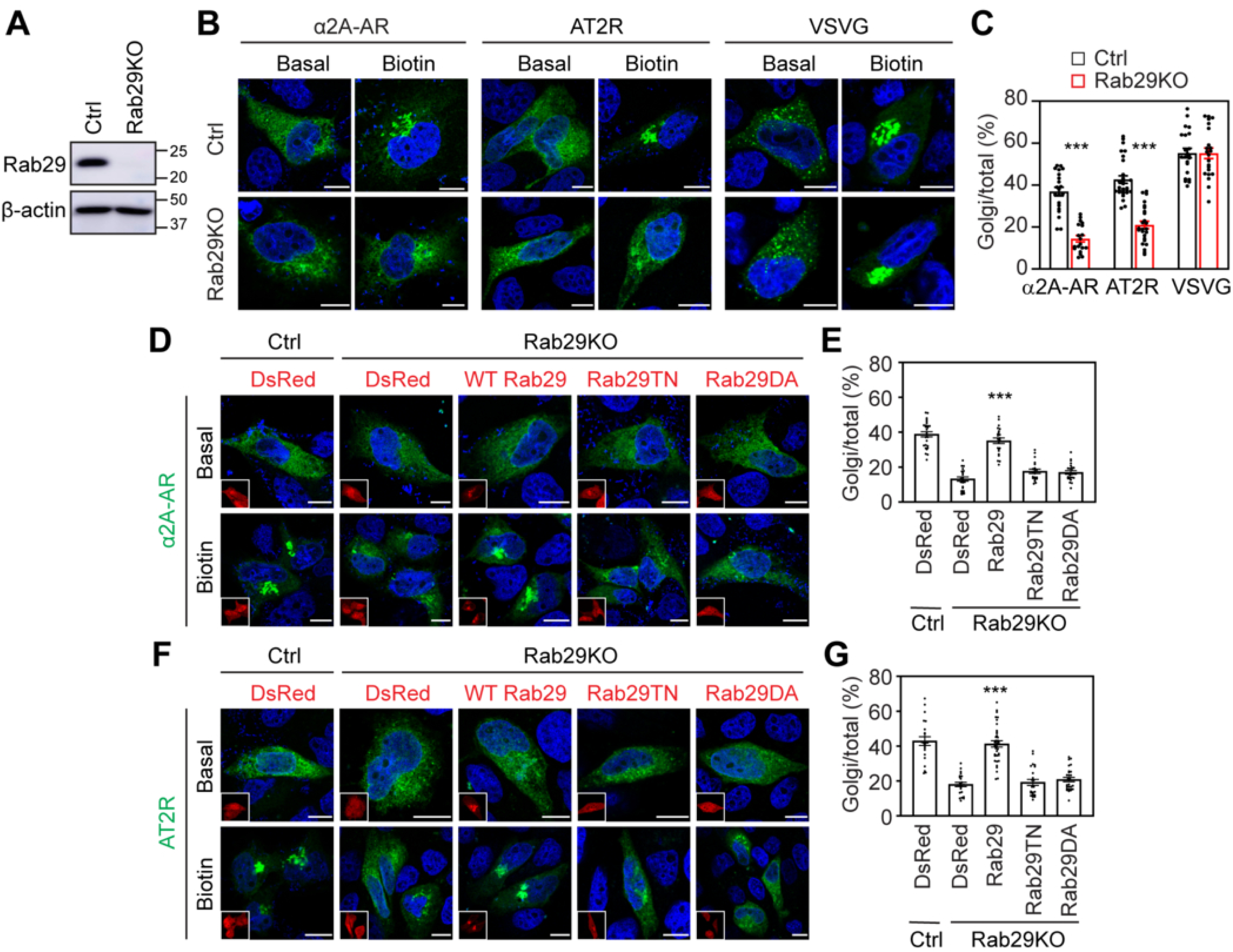
Rab29 KO attenuates the ER-to-Golgi transport of GPCRs in HEK293 cells. **(A)** Rab29 expression in control and Rab29 KO HEK293 cells. **(B)** Rab29 KO inhibits the ER-to-Golgi transport of α_2A_-AR and AT2R, but not VSVG. **(C)** Quantitative data shown in (B). **(D)** and **(F)** Rescue of the ER-to-Golgi transport of α_2A_-AR (D) and AT2R (F) in Rab29 KO cells by Rab29. HEK293 cells were transfected with RUSH α_2A_-AR or AT2R together with DsRed or CRSIPR-Cas9- resistant DsRed-Rab29 and then treated with biotin for 15 min. Insets show the expression of DsRed or DsRed-Rab29. **(E)** and **(G)** Quantitative data shown in (D) and (F), respectively. The quantitative data are means ± SE; n = 20-29 cells in (C), 20-27 cells in (E), and 20-36 cells in (G), each from 3-5 repeats. ***p < 0.001 versus control in (C) and versus DsRed, Rab29TN, and Rab29DA in Rab29 KO cells in (E) and (G). Scale bars,10 µm.

**Figure 3 – figure supplement 1.**
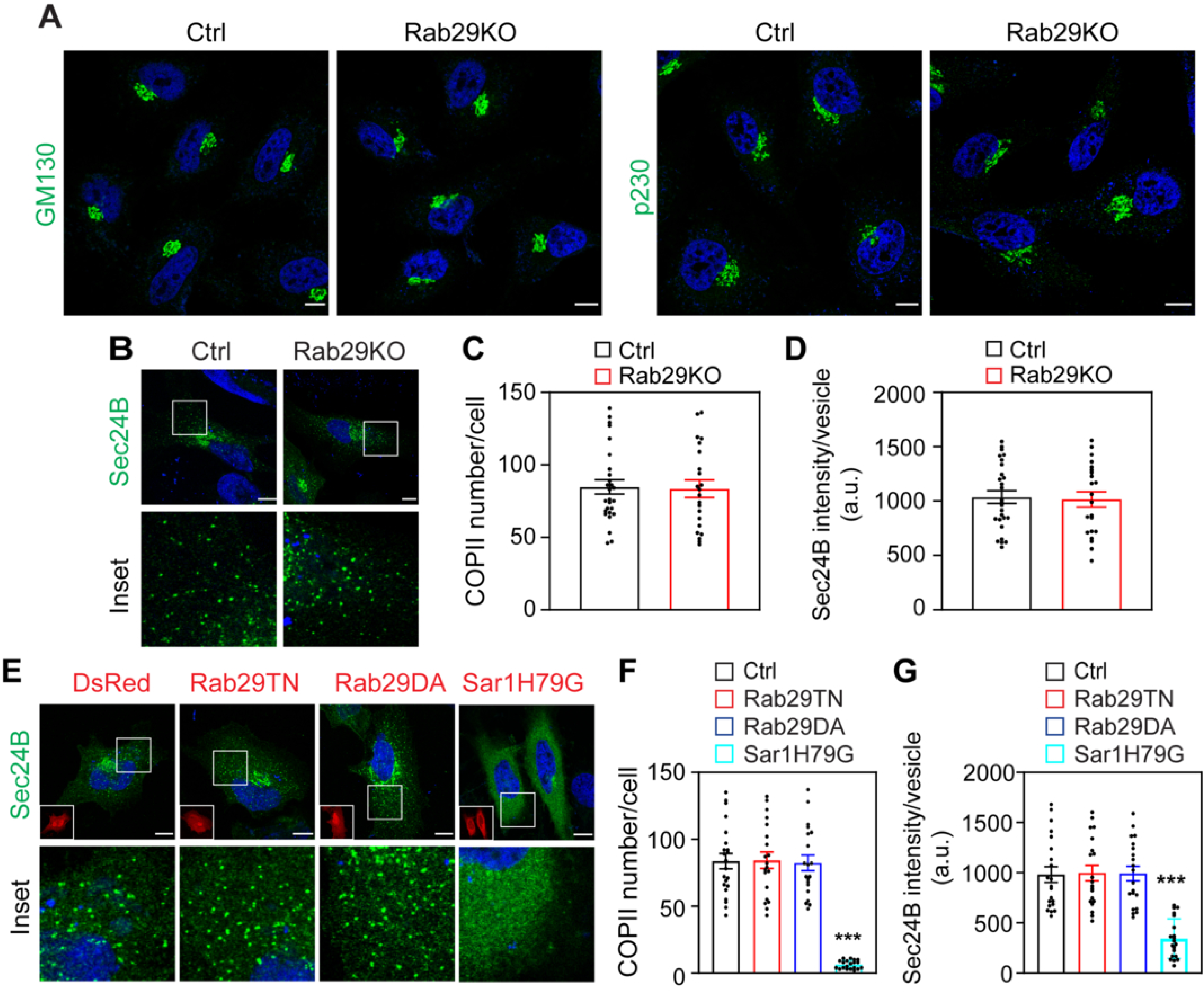
Effects of Rab29 KO and its inactive mutants on Golgi structure and the formation of COPII vesicles in HeLa cells. **(A)** Representative images showing the overall Golgi structure after staining with GM130 and p230 antibodies in control and Rab29 KO cells. (B) and **(E)** Effects of Rab29 KO (B) and Rab29 and Sar1 mutants (E) on the formation of COPII vesicles using GFP-Sec24B as a marker. **(C)** and **(F)** Quantitative data showing the number of COPII vesicles per cell. **(D)** and **(G)** Quantitative data showing the average Sec24B expression in COPII vesicles. The quantitative data are means ± SE; n = 20-27 cells from 3-5 repeats. ***p < 0.001 versus control. Scale bars,10 mm. Insets on the bottom left of the upper panel in (E) show the expression of DsRed and DsRed-tagged Rab29 and Sar1 mutants.

**Figure 3 – figure supplement 2.**
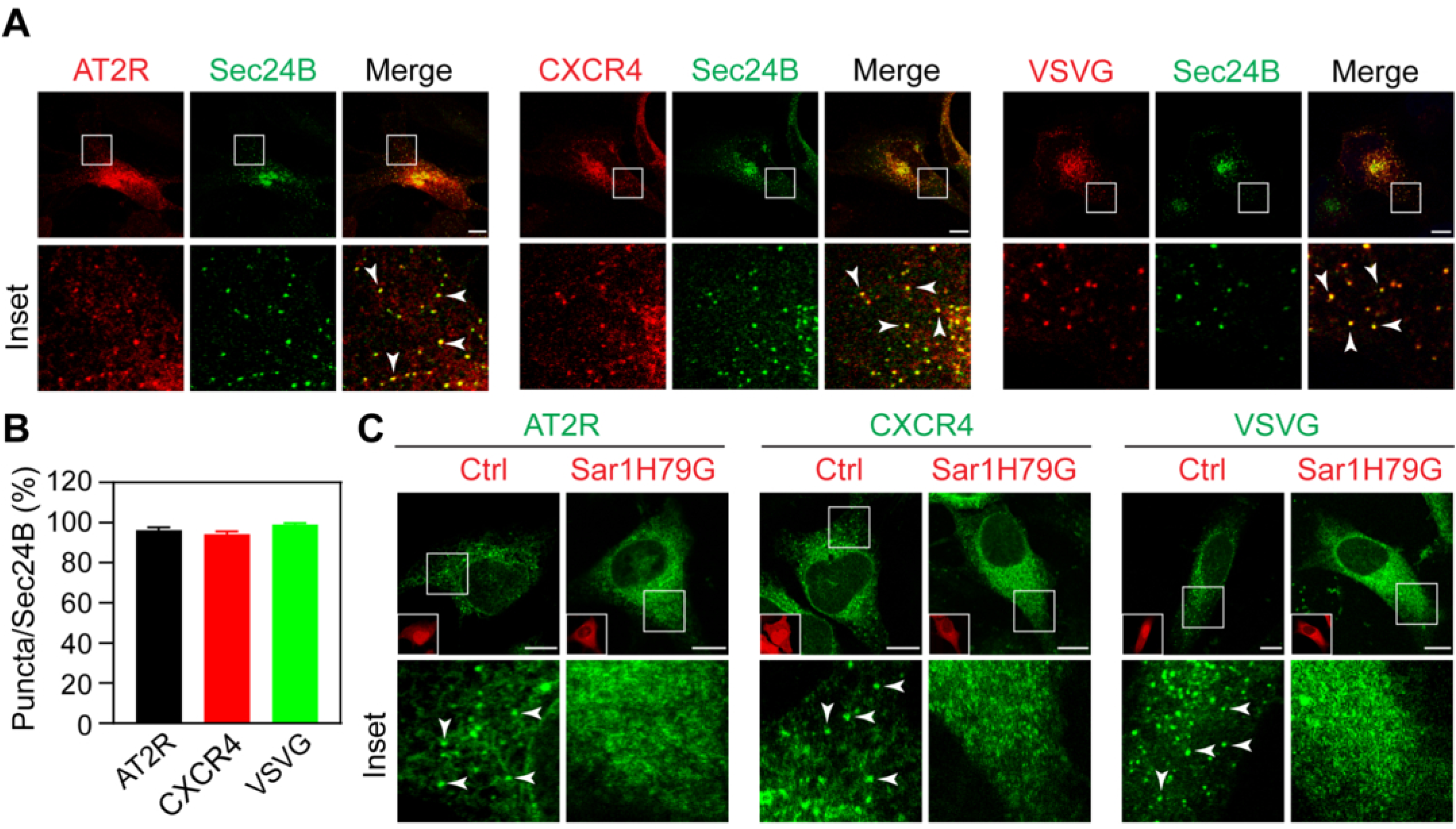
Expression of RUSH AT2R, CXCR4, and VSVG at COPII vesicles. **(A)** Colocalization of RUSH AT2R, CXCR4, and VSVG with Sec24B as a COPII marker. HeLa cells were transfected with mCherry-tagged RUSH cargoes and GFP-Sec24B. **(B)** Quantification of individual cargoes in COPII vesicles. The data shown are the percentage of cargo-carrying punctate structures that contained Sec24B. **(C)** Sar1H79G inhibits the expression of RUSH AT2R, CXCR4, and VSVG in COPII vesicles. Insets on the bottom left in (C) show the expression of DsRed or DsRed- Sar1H79G. Arrows indicate cargo concentration in COPII vesicles. Scale bars,10 µm.

**Figure 5 – figure supplement 1.**
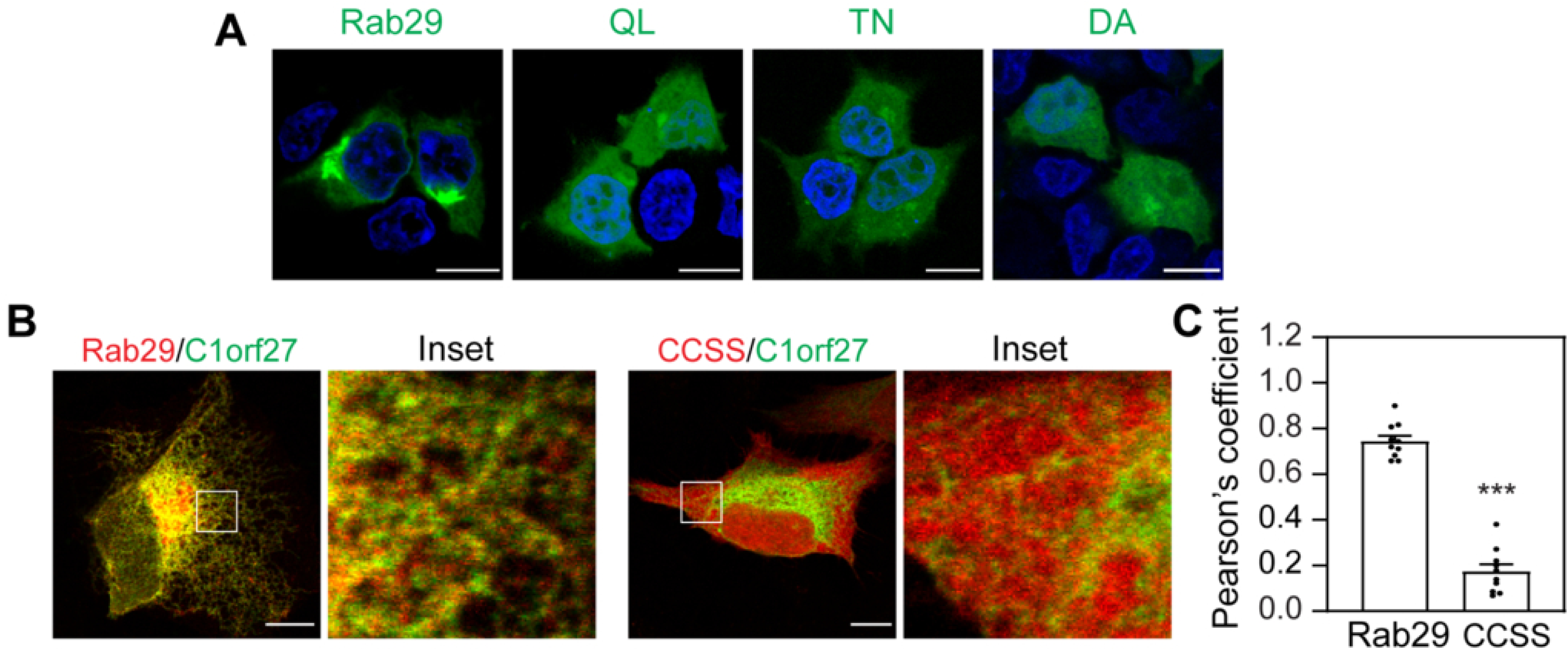
Rab29 localizes to the ER in an activation- and geranylgeranylation-dependent fashion. **(A)** Subcellular distribution of GFP-tagged Rab29 and its QL, TN, and DA mutants in HEK293 cells. **(B)** Effect of mutating the C-terminal CC motif to SS on the ER localization of Rab29. HeLa cells were transfected with GFP-C1orf27 and DsRed-Rab29 or its CCSS mutant. **(C)** Quantification of Pearson’s correlation coefficient between Rab29 and C1orf27. The quantitative data are means ± SE; n = 10 cells from 4 experiments in (C). ***p < 0.001 versus Rab29. Scale bars, 10 µm.

**Figure 6 – figure supplement 1.**
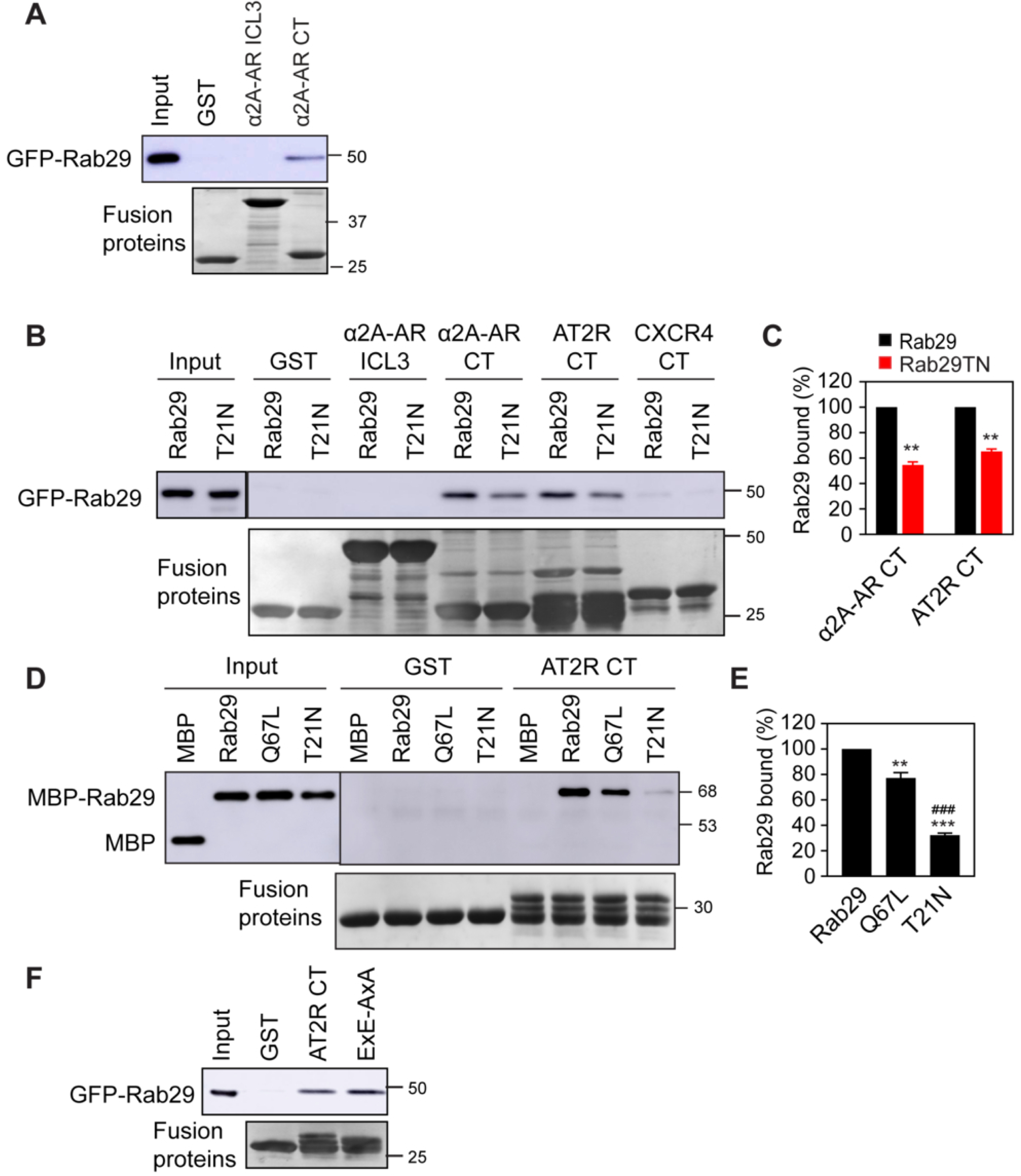
Interaction of intracellular domains of α_2A_-AR, AT2R, and CXCR4 with Rab29. **(A)** Interaction of the ICL3 and CT of α_2A_-AR with Rab29 in GST pulldown assays. **(B)** Interaction of the intracellular domains of α_2A_-AR, AT2R, and CXCR4 with Rab29 and its TN mutant. Individual receptor domains were generated as GST fusion proteins and incubated with cell lysates expressing GFP-Rab29. **(C)** Quantitative data shown in (B). **(D)** Interaction of GST-AT2R CT with purified MBP-Rab29 and its QL and TN mutants. **(E)** Quantitative data shown in (D). **(F)** Interaction of the AT2R CT and its ExE-AxA mutant with GFP-Rab29. Similar results were obtained in at least 3 independent experiments for each panel. Input lanes contain 2% of the total protein used in the experiments. The quantitative data are means ± SE; n = 3 separate experiments. *p < 0.05, **p < 0.01 and ***p < 0.001 versus Rab29; ^###^p < 0.001 versus QL.

## References

Alessi, D.R., and S.R. Pfeffer. 2024. Leucine-Rich Repeat Kinases. Annu Rev Biochem. 93:261–287.

Allan, B.B., B.D. Moyer, and W.E. Balch. 2000. Rab1 recruitment of p115 into a cis-SNARE complex: programming budding COPII vesicles for fusion. Science. 289:444–448.

Appenzeller, C., H. Andersson, F. Kappeler, and H.P. Hauri. 1999. The lectin ERGIC-53 is a cargo transport receptor for glycoproteins. Nat Cell Biol. 1:330–334.

Assoumou, K., S. Papadogkonaki, I. Muneta-Arrate, and M. Stoeber. 2025. Mechanisms governing GPCR anterograde transport. FEBS Lett. 599:2420–2438.

Bajaj Pahuja, K., J. Wang, A. Blagoveshchenskaya, L. Lim, M.S. Madhusudhan, P. Mayinger, and R. Schekman. 2015. Phosphoregulatory protein 14-3-3 facilitates SAC1 transport from the endoplasmic reticulum. Proc Natl Acad Sci U S A. 112:E3199–3206.

Barlowe, C., and A. Helenius. 2016. Cargo capture and bulk flow in the early secretory pathway. Annu Rev Cell Dev Biol. 32:197–222.

Beilina, A., L. Bonet-Ponce, R. Kumaran, J.J. Kordich, M. Ishida, A. Mamais, A. Kaganovich, S. Saez- Atienzar, D.C. Gershlick, D.A. Roosen, L. Pellegrini, V. Malkov, M.J. Fell, K. Harvey, J.S. Bonifacino, D.J. Moore, and M.R. Cookson. 2020. The Parkinson’s disease protein LRRK2 interacts with the GARP complex to promote retrograde transport to the trans-Golgi network. Cell Rep. 31:107614.

Beilina, A., I.N. Rudenko, A. Kaganovich, L. Civiero, H. Chau, S.K. Kalia, L.V. Kalia, E. Lobbestael, R. Chia, K. Ndukwe, J. Ding, M.A. Nalls, C. International Parkinson’s Disease Genomics, C. North American Brain Expression, M. Olszewski, D.N. Hauser, R. Kumaran, A.M. Lozano, V. Baekelandt, L.E. Greene, J.M. Taymans, E. Greggio, and M.R. Cookson. 2014. Unbiased screen for interactors of leucine-rich repeat kinase 2 supports a common pathway for sporadic and familial Parkinson disease. Proc Natl Acad Sci U S A. 111:2626–2631.

Boncompain, G., S. Divoux, N. Gareil, H. de Forges, A. Lescure, L. Latreche, V. Mercanti, F. Jollivet, G. Raposo, and F. Perez. 2012. Synchronization of secretory protein traffic in populations of cells. Nat Methods. 9:493–498.

Carroll, K.S., J. Hanna, I. Simon, J. Krise, P. Barbero, and S.R. Pfeffer. 2001. Role of Rab9 GTPase in facilitating receptor recruitment by TIP47. Science. 292:1373–1376.

Conn, P.M., A. Ulloa-Aguirre, J. Ito, and J.A. Janovick. 2007. G protein-coupled receptor trafficking in health and disease: lessons learned to prepare for therapeutic mutant rescue in vivo. Pharmacol Rev. 59:225–250.

Di Minin, G., M. Holzner, A. Grison, C.E. Dumeau, W. Chan, A. Monfort, L.A. Jerome-Majewska, H. Roelink, and A. Wutz. 2022. TMED2 binding restricts SMO to the ER and Golgi compartments. PLoS Biol. 20:e3001596.

di Ronza, A., L. Bajaj, J. Sharma, D. Sanagasetti, P. Lotfi, C.J. Adamski, J. Collette, M. Palmieri, A. Amawi, L. Popp, K.T. Chang, M.C. Meschini, H.E. Leung, L. Segatori, A. Simonati, R.N. Sifers, F.M. Santorelli, and M. Sardiello. 2018. CLN8 is an endoplasmic reticulum cargo receptor that regulates lysosome biogenesis. Nat Cell Biol. 20:1370–1377.

Doly, S., H. Shirvani, G. Gäta, F.J. Meye, M.B. Emerit, H. Enslen, L. Achour, L. Pardo-Lopez, S.K. Yang, V. Armand, R. Gardette, B. Giros, M. Gassmann, B. Bettler, M. Mameli, M. Darmon, and S. Marullo. 2016. GABAB receptor cell-surface export is controlled by an endoplasmic reticulum gatekeeper. Mol Psychiatry. 21:480–490.

Downes, K.W., and G. Zanetti. 2025. Mechanisms of COPII coat assembly and cargo recognition in the secretory pathway. Nat Rev Mol Cell Biol. 26:910–925.

Duvernay, M.T., F. Zhou, and G. Wu. 2004. A conserved motif for the transport of G protein-coupled receptors from the endoplasmic reticulum to the cell surface. J Biol Chem. 279:30741–30750.

Eguchi, T., T. Kuwahara, M. Sakurai, T. Komori, T. Fujimoto, G. Ito, S.I. Yoshimura, A. Harada, M. Fukuda, M. Koike, and T. Iwatsubo. 2018. LRRK2 and its substrate Rab GTPases are sequentially targeted onto stressed lysosomes and maintain their homeostasis. Proc Natl Acad Sci U S A. 115:E9115–E9124.

Emmer, B.T., G.G. Hesketh, E. Kotnik, V.T. Tang, P.J. Lascuna, J. Xiang, A.C. Gingras, X.W. Chen, and D. Ginsburg. 2018. The cargo receptor SURF4 promotes the efficient cellular secretion of PCSK9. Elife. 7.

Farhan, H., I. Raote, F. Campelo, L. Ge, K. Hirschberg, A. Forrester, G. Zanetti, J. Lippincott-Schwartz, J.C. Pastor-Pareja, F. Perez, K. Saito, and V. Malhotra. 2025. Towards a unified framework for the function of endoplasmic reticulum exit sites. Nat Rev Mol Cell Biol. 26:957–969.

Fath, S., J.D. Mancias, X. Bi, and J. Goldberg. 2007. Structure and organization of coat proteins in the COPII cage. Cell. 129:1325–1336.

Gomez, R.C., P. Wawro, P. Lis, D.R. Alessi, and S.R. Pfeffer. 2019. Membrane association but not identity is required for LRRK2 activation and phosphorylation of Rab GTPases. J Cell Biol. 218:4157–4170.

Haas, A.K., S. Yoshimura, D.J. Stephens, C. Preisinger, E. Fuchs, and F.A. Barr. 2007. Analysis of GTPase-activating proteins: Rab1 and Rab43 are key Rabs required to maintain a functional Golgi complex in human cells. J Cell Sci. 120:2997–3010.

Homma, Y., R. Kinoshita, Y. Kuchitsu, P.S. Wawro, S. Marubashi, M.E. Oguchi, M. Ishida, N. Fujita, and M. Fukuda. 2019. Comprehensive knockout analysis of the Rab family GTPases in epithelial cells. J Cell Biol. 218:2035–2050.

Huang, Y., H. Yin, B. Li, Q. Wu, Y. Liu, K. Poljak, J. Maldutyte, X. Tang, M. Wang, Z. Wu, E.A. Miller, L. Jiang, Z.P. Yao, and Y. Guo. 2021. An in vitro vesicle formation assay reveals cargo clients and factors that mediate vesicular trafficking. Proc Natl Acad Sci U S A. 118:e2101287118.

Jang, W., K. Senarath, G. Feinberg, S. Lu, and N.A. Lambert. 2024. Visualization of endogenous G proteins on endosomes and other organelles. Elife. 13:RP97033.

Jones-Tabah, J. 2023. Targeting G protein-coupled receptors in the treatment of Parkinson’s disease. J Mol Biol. 435:167927.

Komori, T., T. Kuwahara, T. Fujimoto, M. Sakurai, I. Koyama-Honda, M. Fukuda, and T. Iwatsubo. 2023. Phosphorylation of Rab29 at Ser185 regulates its localization and role in the lysosomal stress response in concert with LRRK2. J Cell Sci. 136.

Kunselman, J.M., J. Lott, and M.A. Puthenveedu. 2021. Mechanisms of selective G protein-coupled receptor localization and trafficking. Curr Opin Cell Biol. 71:158–165.

Li, C., Y. Fan, T.H. Lan, N.A. Lambert, and G. Wu. 2012. Rab26 modulates the cell surface transport of alpha2-adrenergic receptors from the Golgi. J Biol Chem. 287:42784–42794.

Li, C., Z. Wei, Y. Fan, W. Huang, Y. Su, H. Li, Z. Dong, M. Fukuda, M. Khater, and G. Wu. 2017. The GTPase Rab43 controls the anterograde ER-Golgi trafficking and sorting of GPCRs. Cell Rep. 21:1089–1101.

Liu, S., P.J. Anderson, S. Rajagopal, R.J. Lefkowitz, and H.A. Rockman. 2024. G protein-coupled receptors: A century of research and discovery. Circ Res. 135:174–197.

Liu, Z., N. Bryant, R. Kumaran, A. Beilina, A. Abeliovich, M.R. Cookson, and A.B. West. 2018. LRRK2 phosphorylates membrane-bound Rabs and is activated by GTP-bound Rab7L1 to promote recruitment to the trans-Golgi network. Hum Mol Genet. 27:385–395.

Lorente, J.S., A.V. Sokolov, G. Ferguson, H.B. Schiöth, A.S. Hauser, and D.E. Gloriam. 2025. GPCR drug discovery: new agents, targets and indications. Nat Rev Drug Discov.

MacLeod, D.A., H. Rhinn, T. Kuwahara, A. Zolin, G. Di Paolo, B.D. McCabe, K.S. Marder, L.S. Honig, L.N. Clark, S.A. Small, and A. Abeliovich. 2013. RAB7L1 interacts with LRRK2 to modify intraneuronal protein sorting and Parkinson’s disease risk. Neuron. 77:425–439.

Mancias, J.D., and J. Goldberg. 2008. Structural basis of cargo membrane protein discrimination by the human COPII coat machinery. Embo j. 27:2918–2928.

Miller, E.A., T.H. Beilharz, P.N. Malkus, M.C. Lee, S. Hamamoto, L. Orci, and R. Schekman. 2003. Multiple cargo binding sites on the COPII subunit Sec24p ensure capture of diverse membrane proteins into transport vesicles. Cell. 114:497–509.

Mizuno-Yamasaki, E., F. Rivera-Molina, and P. Novick. 2012. GTPase networks in membrane traffic. Annu Rev Biochem. 81:637–659.

Nagai-Ito, Y., L. Xu, K. Ito, Y. Kajihara, G. Ito, and T. Tomita. 2022. The atypical Rab GTPase associated with Parkinson’s disease, Rab29, is localized to membranes. J Biol Chem. 298:102499.

Nie, C., H. Wang, R. Wang, D. Ginsburg, and X.W. Chen. 2018. Dimeric sorting code for concentrative cargo selection by the COPII coat. Proc Natl Acad Sci U S A. 115:E3155–E3162.

Onnis, A., F. Finetti, L. Patrussi, M. Gottardo, C. Cassioli, S. Spano, and C.T. Baldari. 2015. The small GTPase Rab29 is a common regulator of immune synapse assembly and ciliogenesis. Cell Death Differ. 22:1687–1699.

Petaja-Repo, U.E., M. Hogue, A. Laperriere, P. Walker, and M. Bouvier. 2000. Export from the endoplasmic reticulum represents the limiting step in the maturation and cell surface expression of the human delta opioid receptor. J Biol Chem. 275:13727–13736.

Pfeffer, S.R. 2023. LRRK2 phosphorylation of Rab GTPases in Parkinson’s disease. FEBS Lett. 597:811–818.

Pradhan, G., R. Shrivastva, and S. Mukhopadhyay. 2018. Mycobacterial PknG targets the Rab7l1 signaling pathway to inhibit phagosome-lysosome fusion. J Immunol. 201:1421–1433.

Purlyte, E., H.S. Dhekne, A.R. Sarhan, R. Gomez, P. Lis, M. Wightman, T.N. Martinez, F. Tonelli, S.R. Pfeffer, and D.R. Alessi. 2018. Rab29 activation of the Parkinson’s disease-associated LRRK2 kinase. Embo j. 37:1–18.

Qiu, Z.P., Z.C. Lin, A. Hu, Y.B. Liu, W.E. Zeng, X. Zhao, X.J. Shi, J. Luo, and B.L. Song. 2023. GRAMD1/ASTER-mediated cholesterol transport promotes Smoothened cholesterylation at the endoplasmic reticulum. Embo j. 42:e111513.

Radoux-Mergault, A., L. Oberhauser, S. Aureli, F.L. Gervasio, and M. Stoeber. 2023. Subcellular location defines GPCR signal transduction. Sci Adv. 9:eadf6059.

Sauvageau, E., M.D. Rochdi, M. Oueslati, F.F. Hamdan, Y. Percherancier, J.C. Simpson, R. Pepperkok, and M. Bouvier. 2014. CNIH4 interacts with newly synthesized GPCR and controls their export from the endoplasmic reticulum. Traffic. 15:383–400.

Shiwarski, D.J., A. Tipton, M.D. Giraldo, B.F. Schmidt, M.S. Gold, A.A. Pradhan, and M.A. Puthenveedu. 2017. A PTEN-regulated checkpoint controls surface delivery of δ opioid receptors. J Neurosci. 37:3741–3752.

Simón-Sánchez, J., C. Schulte, J.M. Bras, M. Sharma, J.R. Gibbs, D. Berg, C. Paisan-Ruiz, P. Lichtner, S.W. Scholz, D.G. Hernandez, R. Krüger, M. Federoff, C. Klein, A. Goate, J. Perlmutter, M. Bonin, M.A. Nalls, T. Illig, C. Gieger, H. Houlden, M. Steffens, M.S. Okun, B.A. Racette, M.R. Cookson, K.D. Foote, H.H. Fernandez, B.J. Traynor, S. Schreiber, S. Arepalli, R. Zonozi, K. Gwinn, M. van der Brug, G. Lopez, S.J. Chanock, A. Schatzkin, Y. Park, A. Hollenbeck, J. Gao, X. Huang, N.W. Wood, D. Lorenz, G. Deuschl, H. Chen, O. Riess, J.A. Hardy, A.B. Singleton, and T. Gasser. 2009. Genome- wide association study reveals genetic risk underlying Parkinson’s disease. Nat Genet. 41:1308–1312.

Slavin, I., I.A. García, P. Monetta, H. Martinez, N. Romero, and C. Alvarez. 2011. Role of Rab1b in COPII dynamics and function. Eur J Cell Biol. 90:301–311.

Steckelings, U.M., R.E. Widdop, E.D. Sturrock, L. Lubbe, T. Hussain, E. Kaschina, T. Unger, A. Hallberg, R.M. Carey, and C. Sumners. 2022. The angiotensin AT(2) receptor: from a binding site to a novel therapeutic target. Pharmacol Rev. 74:1051–1135.

Steger, M., F. Diez, H.S. Dhekne, P. Lis, R.S. Nirujogi, O. Karayel, F. Tonelli, T.N. Martinez, E. Lorentzen, S.R. Pfeffer, D.R. Alessi, and M. Mann. 2017. Systematic proteomic analysis of LRRK2- mediated Rab GTPase phosphorylation establishes a connection to ciliogenesis. Elife. 6:e31012.

Stenmark, H. 2009. Rab GTPases as coordinators of vesicle traffic. Nat Rev Mol Cell Biol. 10:513–525.

Tang, V.T., and D. Ginsburg. 2023. Cargo selection in endoplasmic reticulum-to-Golgi transport and relevant diseases. J Clin Invest. 133:e163838.

Tang, X., R. Chen, V.S.D. Mesias, T. Wang, Y. Wang, K. Poljak, X. Fan, H. Miao, J. Hu, L. Zhang, J. Huang, S. Yao, E.A. Miller, and Y. Guo. 2022. A SURF4-to-proteoglycan relay mechanism that mediates the sorting and secretion of a tagged variant of sonic hedgehog. Proc Natl Acad Sci U S A. 119:e2113991119.

Tucci, A., M.A. Nalls, H. Houlden, T. Revesz, A.B. Singleton, N.W. Wood, J. Hardy, and C. Paisán-Ruiz. 2010. Genetic variability at the PARK16 locus. Eur J Hum Genet. 18:1356–1359.

Vides, E.G., A. Adhikari, C.Y. Chiang, P. Lis, E. Purlyte, C. Limouse, J.L. Shumate, E. Spinola-Lasso, H.S. Dhekne, D.R. Alessi, and S.R. Pfeffer. 2022. A feed-forward pathway drives LRRK2 kinase membrane recruitment and activation. Elife. 11:e79771.

von Zastrow, M., and A. Sorkin. 2021. Mechanisms for Regulating and Organizing Receptor Signaling by Endocytosis. Annu Rev Biochem. 90:709–737.

Wang, L., H. Liu, X. Zhang, E. Song, Y. Wang, T. Xu, and Z. Li. 2021a. WFS1 functions in ER export of vesicular cargo proteins in pancreatic β-cells. Nat Commun. 12:6996.

Wang, S., Z. Ma, X. Xu, Z. Wang, L. Sun, Y. Zhou, X. Lin, W. Hong, and T. Wang. 2014. A role of Rab29 in the integrity of the trans-Golgi network and retrograde trafficking of mannose-6-phosphate receptor. PLoS One. 9:e96242.

Wang, X., H. Wang, B. Xu, D. Huang, C. Nie, L. Pu, G.J.M. Zajac, H. Yan, J. Zhao, F. Shi, B.T. Emmer, J. Lu, R. Wang, X. Dong, J. Dai, W. Zhou, C. Wang, G. Gao, Y. Wang, C. Willer, X. Lu, Y. Zhu, and X.W. Chen. 2021b. Receptor-mediated ER export of lipoproteins controls lipid homeostasis in mice and humans. Cell Metab. 33:350–366.e357.

Wei, Z., M. Zhang, C. Li, W. Huang, Y. Fan, J. Guo, M. Khater, M. Fukuda, Z. Dong, G. Hu, and G. Wu. 2019. Specific TBC domain-containing proteins control the ER-Golgi-plasma membrane trafficking of GPCRs. Cell Rep. 28:554–566 e554.

Xu, X., Y. Fang, L. Qiu, W. Huang, H. Li, and G. Wu. 2026. The UFSP2-C1orf27 complex positions nascent GPCRs to the ufmylation system for ER export control. iScience. 29:115707.

Xu, X., W. Huang, C.N. Bryant, Z. Dong, H. Li, and G. Wu. 2024a. The ufmylation cascade controls COPII recruitment, anterograde transport, and sorting of nascent GPCRs at ER. Sci Adv. 10:eadm9216.

Xu, X., N.A. Lambert, and G. Wu. 2023. Sequence-directed concentration of G protein-coupled receptors in COPII vesicles. iScience. 26:107969.

Xu, X., L. Qiu, M. Zhang, and G. Wu. 2024b. Segregation of nascent GPCRs in the ER-to-Golgi transport by CCHCR1 via direct interaction. J Cell Sci. 137.

Xu, X., and G. Wu. 2022. Human C1orf27 protein interacts with alpha(2A)-adrenergic receptor and regulates its anterograde transport. J Biol Chem. 298:102021.

Zhang, H., A. Luginina, A. Mishin, M. Baidya, A.K. Shukla, and V. Cherezov. 2021. Structural insights into ligand recognition and activation of angiotensin receptors. Trends Pharmacol Sci. 42:577–587.

Zhang, M., and G. Wu. 2019. Mechanisms of the anterograde trafficking of GPCRs: Regulation of AT1R transport by interacting proteins and motifs. Traffic. 20:110–120.

Zhang, P., and R. Schekman. 2016. Distinct stages in the recognition, sorting, and packaging of proTGFα into COPII-coated transport vesicles. Mol Biol Cell. 27:1938–1947.

Zhang, X., C. Dong, Q.J. Wu, W.E. Balch, and G. Wu. 2011. Di-acidic motifs in the membrane-distal C termini modulate the transport of angiotensin II receptors from the endoplasmic reticulum to the cell surface. J Biol Chem. 286:20525–20535.

Zhu, H., F. Tonelli, M. Turk, A. Prescott, D.R. Alessi, and J. Sun. 2023. Rab29-dependent asymmetrical activation of leucine-rich repeat kinase 2. Science. 382:1404–1411.

